# Basal forebrain volume is associated with cortical amyloid burden in cognitively unimpaired older adults at varying genetic risk for Alzheimer’s disease

**DOI:** 10.1101/2025.05.28.656557

**Authors:** Gawon Cho, Kaitlin Kelly, Takuya Toyonaga, Richard E Carson, Brienne Miner, Joel Gelernter, Pradeep Varma, Christopher H. van Dyck, Adam P. Mecca, Ryan O’Dell

**Affiliations:** Department of Internal Medicine, Yale School of Medicine, New Haven, CT, USA; Yale Alzheimer’s Disease Research Unit, Yale School of Medicine, New Haven, CT, USA; Geisinger Commonwealth School of Medicine, Scranton, PA, USA; Department of Radiology and Biomedical Imaging, Yale University School of Medicine, New Haven, CT, USA; Department of Psychiatry, Yale School of Medicine, New Haven, CT, USA; Veterans Affairs Connecticut Healthcare System, West Haven, CT, USA; Department of Neuroscience, Yale University School of Medicine, New Haven, CT, USA; Department of Genetics, Yale University School of Medicine, New Haven, CT, USA; Department of Neurology, Yale University School of Medicine, New Haven, CT, USA

## Abstract

**Background:** In mild cognitive impairment and dementia due to Alzheimer’s disease (AD), postmortem and *in vivo* neuroimaging studies have demonstrated significant neuronal loss in the basal forebrain cholinergic system (BFCS), which provides the primary cholinergic input to the cerebral cortex. Within this region, atrophy is most prominent in the nucleus basalis of Meynert (nbM), a group of posteriorly clustered magnocellular neurons in the BFCS. However, less is known surrounding the relationship between amyloid deposition, BFCS atrophy, and medial temporal lobe (MTL) volume loss in the preclinical stages of AD. The current study investigates the relationship between sub-structural BFCS volume and cortical Aβ burden in cognitively unimpaired middle-aged individuals at varying genetic risk for AD.

**Methods:** Cognitively unimpaired participants aged 50-65 with a first-degree family history for AD were genetically screened to select three groups: *APOE* genotype ε4ε4 (n=15), ε3ε4 (n=15), and ε3ε3 (n=15), matched for age and sex. Participants underwent imaging with [^11^C]PiB PET and structural 3T MRI. Distribution volumes ratios (*DVR*) with a whole cerebellum reference region were calculated for [^11^C]PiB PET analyses. BFCS sub-structural volumes were obtained from the SPM8 Anatomy Toolbox (Cholinergic nuclei [Ch] 1-3, Ch4). MTL subregional volumes (entorhinal cortex, hippocampus, amygdala, parahippocampal gyrus) were extracted using Freesurfer.

**Results:** BFCS amyloid burden was highest among *APOE* ε4 homozygotes (Ch1-3, F(2, 42)=3.26, *P*=0.048; Ch4, F(2, 42)=3.82, *P*= 0.03). Ch4 (nbM), but not Ch1-3 volume, was found to be inversely associated with global Aβ burden (Pearson *r*=-0.40, *P*=0.007). MTL subregional volumes were not associated with global Aβ burden in the pooled sample. Exploratory analyses in groups stratified by amyloid positivity demonstrated reduced Ch4 volume (*P*=0.032) and significant inverse associations between Ch4 volume and amyloid burden (Pearson *r* = −0.70, *P*=0.02) in Aβ+ participants.

**Conclusions:** We observed nbM (Ch4), but not MTL volume, to be significantly inversely associated with cortical amyloid burden in cognitively unimpaired, Aβ+, middle-aged adults at varying genetic risk for AD. These findings provide further *in vivo* evidence suggesting that nbM atrophy is an early structural correlate of AD pathogenesis, potentially preceding MTL atrophy.

## 1. INTRODUCTION

Alzheimer’s disease (AD) is a neurodegenerative disorder that causes progressive cognitive and functional impairment, and ultimately, death. In addition to the accumulation of pathologic extracellular amyloid-β peptide (Aβ) and intracellular neurofibrillary tangles of hyperphosphorylated tau, AD pathology is characterized by neurodegeneration, as reflected by synaptic loss^1–3^ and progressive volume reduction.^4^ Approximately 7 million U.S. adults have been estimated to be living with AD.^5^ Therefore, the characterization of early diagnostic biomarkers of AD pathogenesis is critical, as it can facilitate the identification of high-risk individuals and the initiation of preventive interventions.

The basal forebrain cholinergic system (BFCS) is a major source of cholinergic input to the neocortex, hippocampus, and amygdala. It can be divided into anterior (including the medial septal nucleus and vertical and horizontal limbs of the diagonal band of Broca, corresponding to Cholinergic nuclei 1 [Ch1], Ch2, and Ch3, respectively) and posterior (including nucleus basalis of Meynert (nbM), corresponding to Ch4) substructures.^6,7^ There is a wealth of evidence demonstrating cholinergic deficits in AD^8–15^, with more robust volume loss observed in the nbM (compared to Ch1-3) in participants with mild cognitive impairment (MCI),^16–18^ suggesting nbM atrophy as an emerging structural biomarker in prodromal AD.

AD is now conceptualized as a continuum, beginning with a ‘preclinical’ or pre-symptomatic stage, characterized by an absence of objective cognitive deficits despite the presence of accumulating AD pathology.^19–21^ Emerging evidence suggests that atrophy in the BFCS (particularly in the nbM) may serve as an early biomarker of AD pathogenesis during this pre-symptomatic phase, potentially preceding volume loss in medial temporal lobe (MTL) subregions consistently reported to be affected in early AD, including the entorhinal cortex and hippocampus.^22–24^ One early study demonstrated a significant correlation between cortical Aβ burden and posterior basal forebrain volume in cognitively unimpaired older adults with elevated Aβ burden.^25^ A more recent publication utilizing fluid biomarkers of Aβ and tau found significant nbM atrophy in cognitively unimpaired individuals who were tau-positive or met criteria for preclinical AD (amyloid- and tau-positive), while Ch1-3 and hippocampal atrophy was less pronounced until the prodromal stage of AD.^26^

Despite these advances, most prior studies assessed Aβ burden using fluid biomarkers, which lack regional specificity. Although one earlier study incorporated Aβ PET imaging,^25^ it focused on older adults, leaving a critical gap in understanding whether the association between core AD pathology and BFCS atrophy is present in middle-aged populations at risk of AD. Furthermore, no prior studies have directly assessed Aβ burden *within* the BFCS itself, a crucial step toward clarifying the mechanisms driving basal forebrain atrophy. In addition, prior work has not accounted for *APOE* genotype, the strongest genetic risk factor for AD, in evaluating the link between BFCS degeneration and AD pathology. Given the essential role of cholinergic input in neuromodulation,^27^ degeneration of basal forebrain substructures may result in functional deficits within their projection areas, potentially contributing to subsequent neurodegeneration. Therefore, in the current study, we used structural MRI and [^11^C]PiB PET to elucidate further the relationship between global Aβ burden and BFCS volumes in a cohort of cognitively unimpaired middle-aged individuals at three levels of *APOE* risk for AD. We hypothesized that global amyloid burden would be inversely associated with BFCS volume within the nbM (Ch4) but not more anterior regions (Ch1-3). We also investigated the relationship between global amyloid burden and the volumes of MTL subregions, including the entorhinal cortex and hippocampus. Finally, we explored the relationship between BFCS volume and the volumes of its primary projection areas.

## 2. METHODS

### 2.1. Study design and recruitment

The study sample consisted of participants aged 50-65 years with normal cognition and a positive family history of AD (in at least one first degree relative) and was identical to that described in our previous study.^28^ Potentially eligible participants underwent a screening for genetic evaluation, which included a medical history questionnaire and *APOE* genotyping, described previously.^28^ Participants were selected from three APOE genotype groups (ε3ε3, ε3ε4, ε4ε4) and matched for age and sex, and underwent a screening diagnostic evaluation for eligibility as previously described.^28^ Briefly, participants were excluded if they met clinical criteria for MCI^29^ or dementia due to AD^30^, as evidenced by either a CDR > 0^31^ or abnormal memory function (scoring below 1.5 standard deviations of education adjusted cutoff on Logical Memory II^32^or a score below 27 on MMSE).^33^ No subjects with *APOE* ε2-containing genotypes, which are less common, were included. All participants provided written informed consent as approved by the Yale University Human Investigation Committee prior to participating in the study.

### 2.2. Magnetic resonance imaging (MRI) and defining regions of interest (ROI)

Structural MRI was conducted on a 3T Trio Scanner (Siemens Medical Systems, Erlangen, Germany) with a circularly polarized head coil. The MR acquisition involved a T1-weighted sagittal 3D-magnetization prepared rapid gradient echo-fast spoiled gradient recalled echo (3D-MPRAGE-FSPGR) pulse sequence with an inversion-recovery preparation of 300 msec (echo time [TE] = 3.34 msec, inversion time [TI] = 1100 msec, repetition time [TR] = 2500 msec, flip angle 7°, slice thickness 1.0 mm, 176 slices, matrix 256×256, voxel size=1mm x 1mm x 1mm). Cortical reconstruction and volumetric segmentation were performed on FreeSurfer 6.0 (http://surfer.nmr.mhg.harvard.edu/).^34^ FreeSurfer regions of interest (ROIs), including those in medial temporal lobe (MTL), as defined by the gyral-based cortical parcellation of the Desikan-Killiany atlas,^35^ and larger composite ROIs, defined using a method identical to our previous work,^36^ were applied to each participant’s structural scan in native space to extract regional volumes. BFCS ROIs were extracted from probabilistic cytoarchitectonic maps developed by Zaborszky et al.^37^ using the Statistical Parametric Mapping (SPM, version 12.0) Anatomy Toolbox plugin.^38^ Extracted ROIs included anterior cholinergic nuclei 1-3 (Ch1-3; medial septal and vertical and horizontal limbs of the diagonal band of Broca) and posterior cholinergic nuclei 4 (Ch4; nbM) in both the left and right hemispheres. BFCS volumes were extracted through linear transformation of Ch1-3 and Ch4 masks in MNI template space into each participant’s individual MR space.

### 2.3. Positron emission tomography (PET) and tracer kinetic modeling

Participants underwent a [^11^C]Pittsburgh Compound B (PiB) PET scan with the ECAT HR+ (Siemens) in 3D mode. Dynamic scanning was conducted for 90 minutes following an intravenous injection of 15mCi of [^11^C]PiB. Each scan session produced 63 slices, each with 2.4 mm of separation, and the final image resolution of approximately 6 mm. A transmission scan was used for attenuation correction.

Subsequently, PET images were reconstructed into 27 frames, containing 63 axial slices of 128 x 128 voxels (2.1 x 2.1 x 2.4 mm) after corrections for attenuation, randoms, scatter, and deadtime. Software motion correction was applied to the dynamic PET images using a mutual-information algorithm (FSL 3.2; Analysis Group, FMRIB, Oxford, UK) to perform frame-by-frame registration to a summed image obtained between 0-10 minutes of the dynamic scanning session. The summed, motion-corrected PET image was registered to each participant’s structural MRI in native space. Subsequently, parametric images of non-displaceable binding potential (BP_ND_) were computed with a 2-step simplified reference tissue method (SRTM2) from 0-90 minutes,^39^ using a whole cerebellum as the reference region as previously described.^36,40,41^ Partial volume correction was performed using the iterative Yang algorithm,^42,43^ based on the ROIs defined using each participant’s structural MRI scan. BP_ND_ of each Freesurfer and basal forebrain ROIs was converted to distribution volume ratios (*DVR*), where *DVR* = BPND+1. For basal forebrain ROI BP_ND_ extraction, an AAL gray matter mask (GM) was applied to include only GM voxels. Global cortical Aβ burden was calculated as a weighted average of composite regions commonly affected by Aβ burden in AD, which included prefrontal, lateral temporal, posterior cingulate/precuneus, and lateral parietal ROIs.

### 2.4. Statistical analysis

Group differences in participant characteristics by *APOE* 14 genotype were assessed with the Analysis of Variance (ANOVA) for continuous variables and χ2 tests for categorical variables. For primary analyses examining the association of global Aβ burden with BFCS and medial temporal lobe (MTL) ROI volumes, we calculated Pearson’s correlation coefficients between BFCS or MTL ROI volume and global cortical Aβ DVR. In addition to the primary analyses in the pooled sample, we conducted subgroup analyses exploring the associations of BFCS and MTL ROI volumes with global Aβ burden in Aβ+ and Aβ-groups. Two different, previously published *DVR* thresholds of 1.2 and 1.08 were used to classify individuals into Aβ+ vs. Aβ-groups.^44^ Pearson’s correlation coefficients were calculated for the associations of global Aβ burden with BFCS and MTL ROI volumes in Aβ+ and Aβ-groups. Additionally, two-tailed t-tests were used to compare BFCS and MTL ROI volumes between Aβ+ and Aβ-groups.

Furthermore, we examined the association of basal forebrain substructure volumes with their projection area volumes, as delineated by Mesulam.^45^ Specifically, Pearson’s correlations were used to examine associations between Ch1-3 and the hippocampus and between Ch4 and both the amygdala and all cortical Freesurfer ROIs.

For all primary analyses, global Aβ *DVR* was corrected for partial volume effects and all ROI volumes were normalized using each participant’s estimated intracranial volume (eICV). All t-tests were two-tailed, and all statistical analyses were performed using SPSS version 21.0 (IBM Corp.) and Matlab version R2024a (The MathWorks Inc.).

## 3. RESULTS

### 3.1. Participant characteristics

The study sample consisted of 45 cognitively normal participants selected to be evenly distributed across 3 *APOE* genotype groups: 1]31]3 (n = 15), 1]31]4 (n =15), and 1]41]4 (n =15). Forty-seven percent of the sample were men, the mean age was 59.1 years (range = 50-67), and all individuals identified as White (Table 1). As previously described^28^, individuals in each *APOE* genotype group were matched for age and sex during recruitment, and there were no group differences in sex (Table 1; χ2 = 0.0, *P* = 1.0), age (F = 0.0, *P* = 0.99), education (F = 0.06, *P* = 0.95), or cognitive performance as assessed by MMSE (F = 1.43, *P* = 0. 25)(Table 1). As previously demonstrated,^28^ global cortical Aβ burden also differed by *APOE* 1]4 carrier status, showing an increasing trend with each additional *APOE* 1]4 copy number (Supplementary Figure 1; F = 4.21, *P* = 0.022). Regional Aβ burden in basal forebrain substructures significantly differed by *APOE* 1]4 carrier status as well, with *APOE* 1]4 homozygotes showing the highest burden (Figure 1; bilateral Ch1-3, F(2, 42) = 3.26, *P* = 0.048; bilateral Ch4, F(2, 42) = 3.82, *P* = 0.03). These findings remained significant after partial volume correction (Supplementary Figure 1).

**Figure 1.**
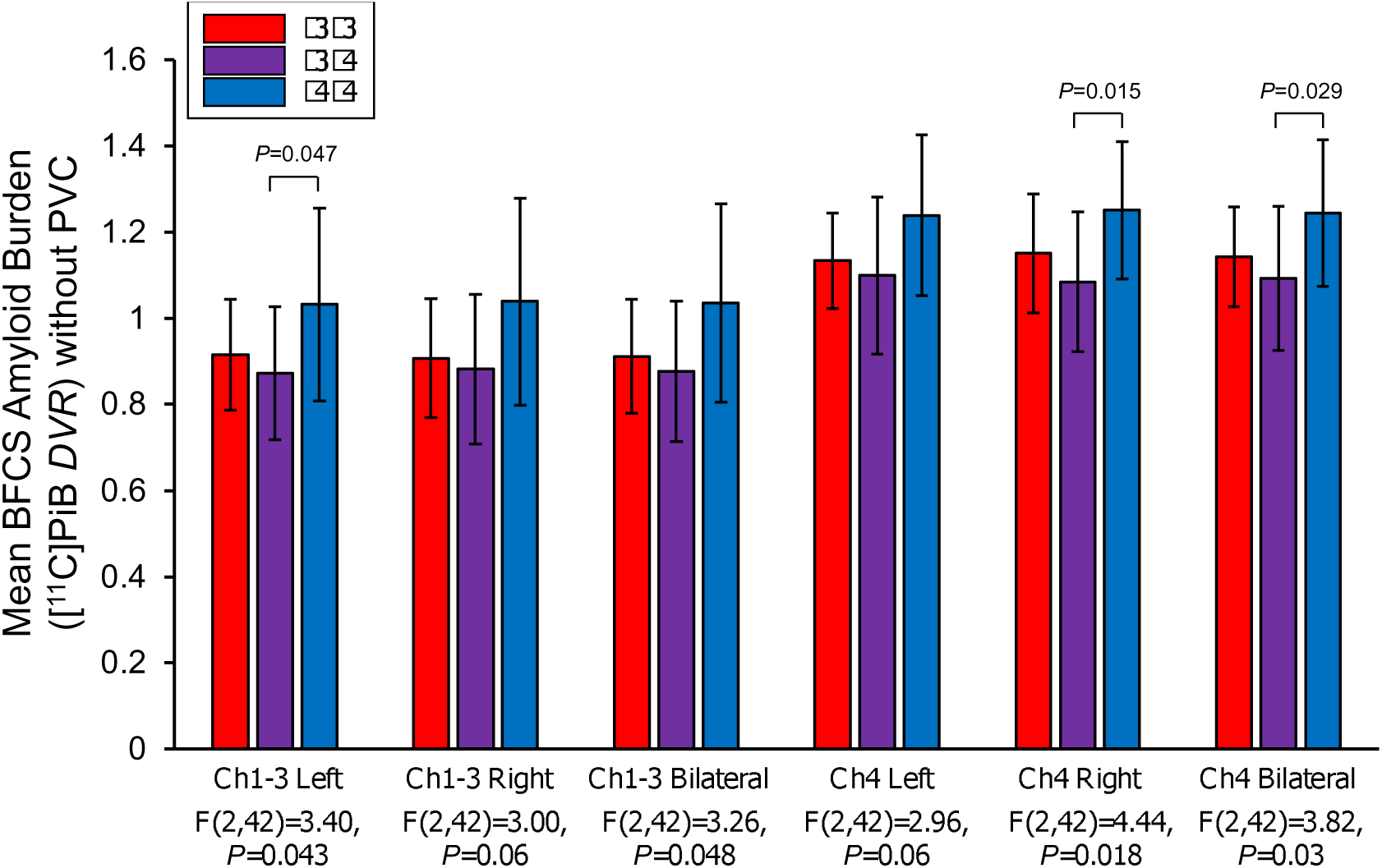
Amyloid burden in basal forebrain substructures by APOE genotype. Group differences in BFCS amyloid burden by APOE genotype were calculated using one-way ANOVA with post-hoc Bonferroni unpaired t-tests. F statistics and P values for each model are depicted below the x-axis and P values from post-hoc unpaired t-tests are depicted above the bar graphs to denote significant group differences. Abbreviations: BFCS, basal forebrain cholinergic system; PiB; Pittsburg compound B; IY-PVC, Iterative Yang partial volume correction; DVR, distribution volume ratio; Ch1-3, cholinergic nuclei 1-3; Ch4, cholinergic nuclei 4.

**Table 1.**
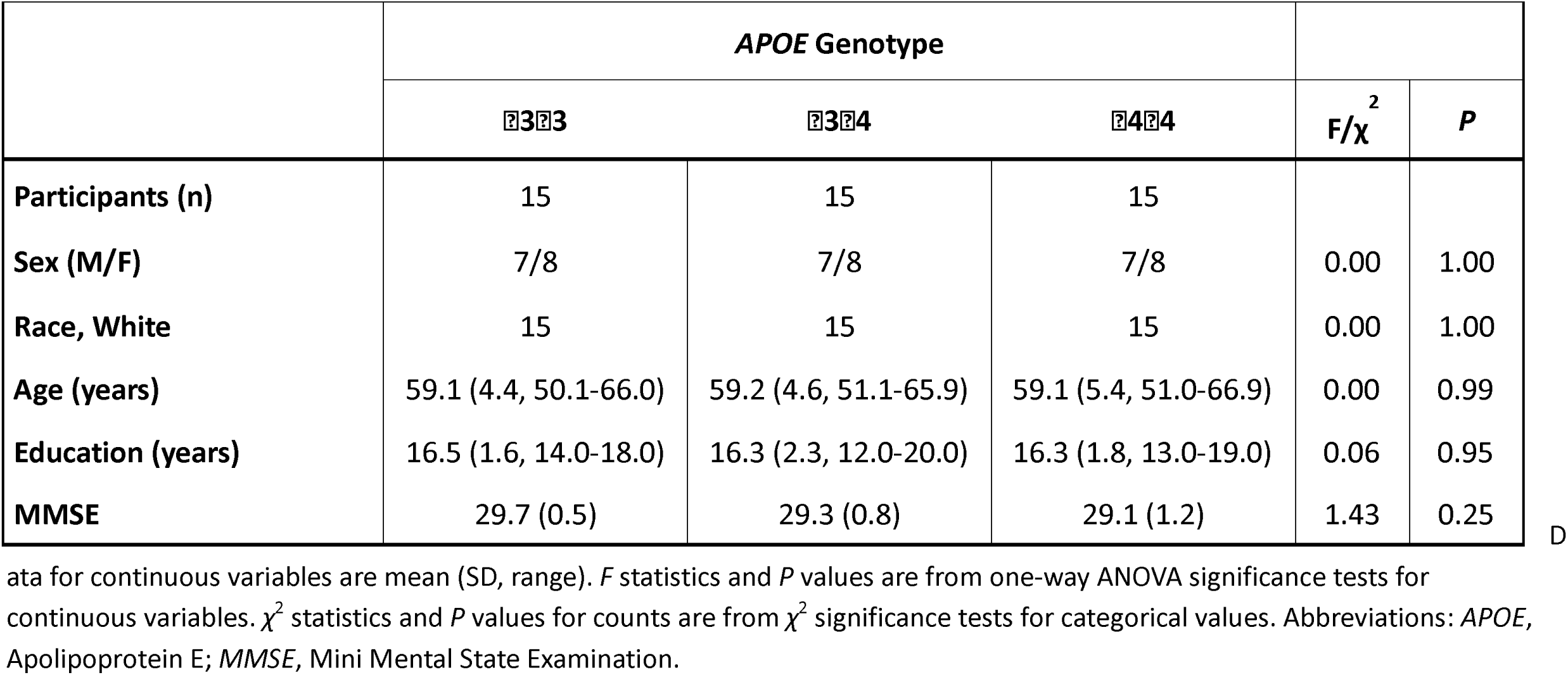
Participant characteristics.

### 3.2. Global and regional Aβ burden and basal forebrain atrophy

For the primary analysis, we investigated the association between global cortical Aβ burden (without partial volume correction) and the volume of each basal forebrain substructure, including the anterior (Ch1-3) and posterior regions (Ch4). Figure 2 depicts scatter plots with zero-order Pearson correlations (*r*). Left, right, and bilateral Ch4 (nbM) volumes were significantly inversely correlated with global Aβ burden, even after covarying for *APOE* genotype (Figure 2D-F; left Ch4, Pearson *r* = −0.36, *P* = 0.02; right Ch4, Pearson *r* = −0.31, *P* = 0.04; bilateral Ch4, Pearson *r* = −0.40, *P* = 0.007). In contrast, anterior basal forebrain volumes (Ch1-3) volumes were not significantly associated with cortical Aβ burden (Figure 2A-C, *P* < 0.05). Results remained virtually identical after partial volume correction (Supplementary Figure 3).

**Figure 2.**
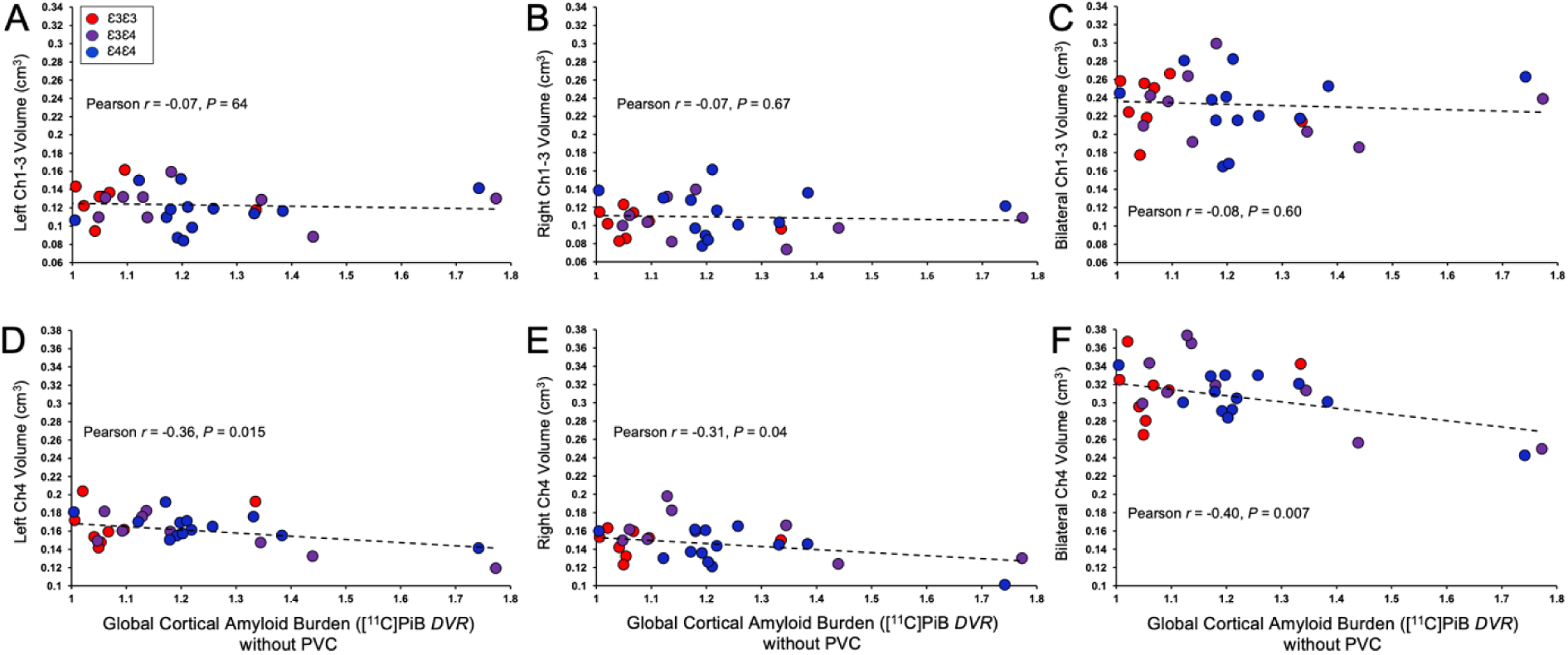
Association between mean global cortical amyloid burden and (A-C) Ch1-3 volume and (D-F) Ch4 volume. Displayed are Pearson r (P) for correlations between global cortical amyloid burden (DVR) and basal forebrain volumes. Abbreviations: PVC, partial volume correction; PiB; Pittsburg compound B; DVR, distribution volume ratio; Ch1-3, cholinergic nuclei 1-3; Ch4, cholinergic nuclei 4.

We further examined variations in the association between basal forebrain substructure volumes and global cortical Aβ burden in Aβ+ and Aβ-participants, using a *DVR* cut-off of 1.2.^44^ Using this cutoff, 11 participants were classified as Aβ+, who were significantly older and more likely to carry one or more copies of *APOE* 1]4 as compared to the 34 Aβ-participants (Supplementary Table 1). Left and bilateral Ch4 volumes were significantly inversely correlated with global cortical Aβ burden in Aβ+ (left Ch4, Pearson *r* = −0.69, *P* = 0.02; bilateral Ch4, Pearson *r* = −0.70, *P* = 0.02), but in not Aβ-participants (Supplementary Figure 4D-F). There were no significant correlations observed between Ch1-3 volume and global cortical Aβ burden in either Aβ+ or Aβ-participants (Supplementary Figure 4A-C). Aβ+ participants were also observed to have significant reductions in Ch4 volume (compared to Aβ-participants), while no group differences were found in Ch1-3 volume (Supplementary Figure 5).

Additional exploratory subgroup analyses using a global cortical Aβ *DVR* threshold of 1.08^44^ to define Aβ positivity also demonstrated a significant inverse correlation between Ch4 volume and global Aβ only among Aβ+ participants (Supplementary Figure 6 and Supplementary Table 2).

Moreover, we examined the association between basal forebrain substructure volumes and each region’s respective Aβ burden using Pearson correlation coefficients (Supplementary Table 3). Regional Aβ burden did not show significant inverse associations with regional volume in any of the basal forebrain substructures. Results remained consistent after partial volume correction.

### 3.3. Global Aβ burden and medial temporal lobe atrophy

Next, we examined the associations between global cortical Aβ burden and MTL ROIs that are known to show signs of atrophy during the prodromal stage of AD, which included the hippocampus, parahippocampal gyrus, entorhinal cortex, and amygdala. In this cohort, non-partial volume corrected global cortical Aβ burden was not correlated with any regional MTL volume, regardless of covarying for *APOE* genotype (Figure 3). Associations remained non-significant after partial volume correction (Supplementary Figure 7). Additionally, there were no significant correlations between global cortical Aβ burden and medial temporal lobe volume in either Aβ+ or Aβ-participants, as defined using a *DVR* cutoff of 1.2 (Supplementary Figure 8). There were no group differences observed in medial temporal lobe volumes between Aβ+ and Aβ-groups (Supplementary Figure 9). Finally, additional sensitivity analyses using a DVR threshold of 1.08 demonstrated comparable results as those based on a *DVR* threshold of 1.2, although a significant inverse correlation between hippocampal volume and global cortical Aβ was found in Aβ-individuals (Supplementary Figure 10).

**Figure 3.**
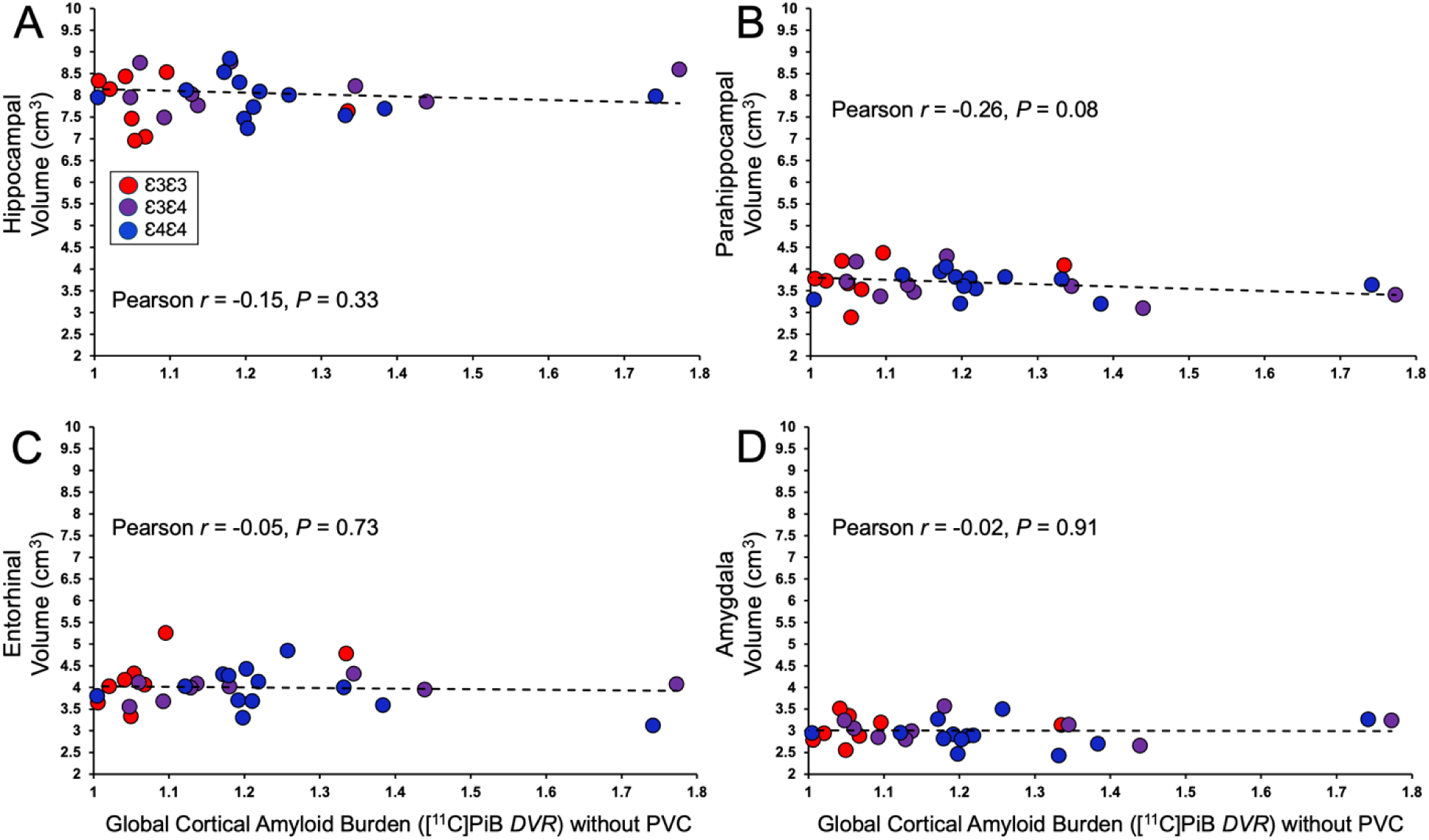
Association between mean global cortical amyloid burden and (A) hippocampal, (B) parahippocampal, (C) entorhinal, and (D) amygdalar volume. Displayed are Pearson r (P) for correlations between global cortical amyloid burden (DVR) and medial temporal lobe volumes. Abbreviations: PVC, partial volume correction; PiB; Pittsburg compound B; DVR, distribution volume ratio.

In addition to the MTL ROIs, we explored the association between all ROIs and global cortical Aβ burden. In the full sample, the volume of bilateral middle temporal gyri showed the strongest inverse association with global cortical Aβ burden (Supplementary Table 4; Pearson *r* = –0.49, P = 0.0006). Middle temporal gyus was the only ROI that showed stronger correlation with cortical Aβ burden than bilateral Ch4. When stratified by Aβ positivity (using a DVR threshold of 1.2^44^), middle temporal gyri volume did not show significant associations with global cortical Aβ burden. These findings remained consistent after applying partial volume correction (Supplementary Table 5).

### 3.4. Relationship between basal forebrain atrophy and volume of primary projection areas

Lastly, we examined the association of basal forebrain substructure volumes with their respective projection area volumes.^45^ Pearson correlations between bilateral Ch1-3 and Ch4 volumes and the volume of each lateralized ROI are shown in Supplementary Table 3 and Figure 4. Anterior basal forebrain volumes (Ch1-3) were not associated with hippocampal volume, the primary projection area for Ch1-3 (left hippocampus, Pearson *r* = 0.19, *P* = 0.21; right hippocampus, Pearson *r* = 0.04, *P =* 0.79; Supplementary Table 3). Additionally, although posterior basal forebrain volume (Ch4, nbM) was observed to be positively correlated with the volumes of a handful of association cortical ROIs (Supplementary Table 6; Figure 4), the observed associations were not broadly distributed across the cerebral cortex, i.e., primary projection area of Ch4.^45^

**Figure 4.**
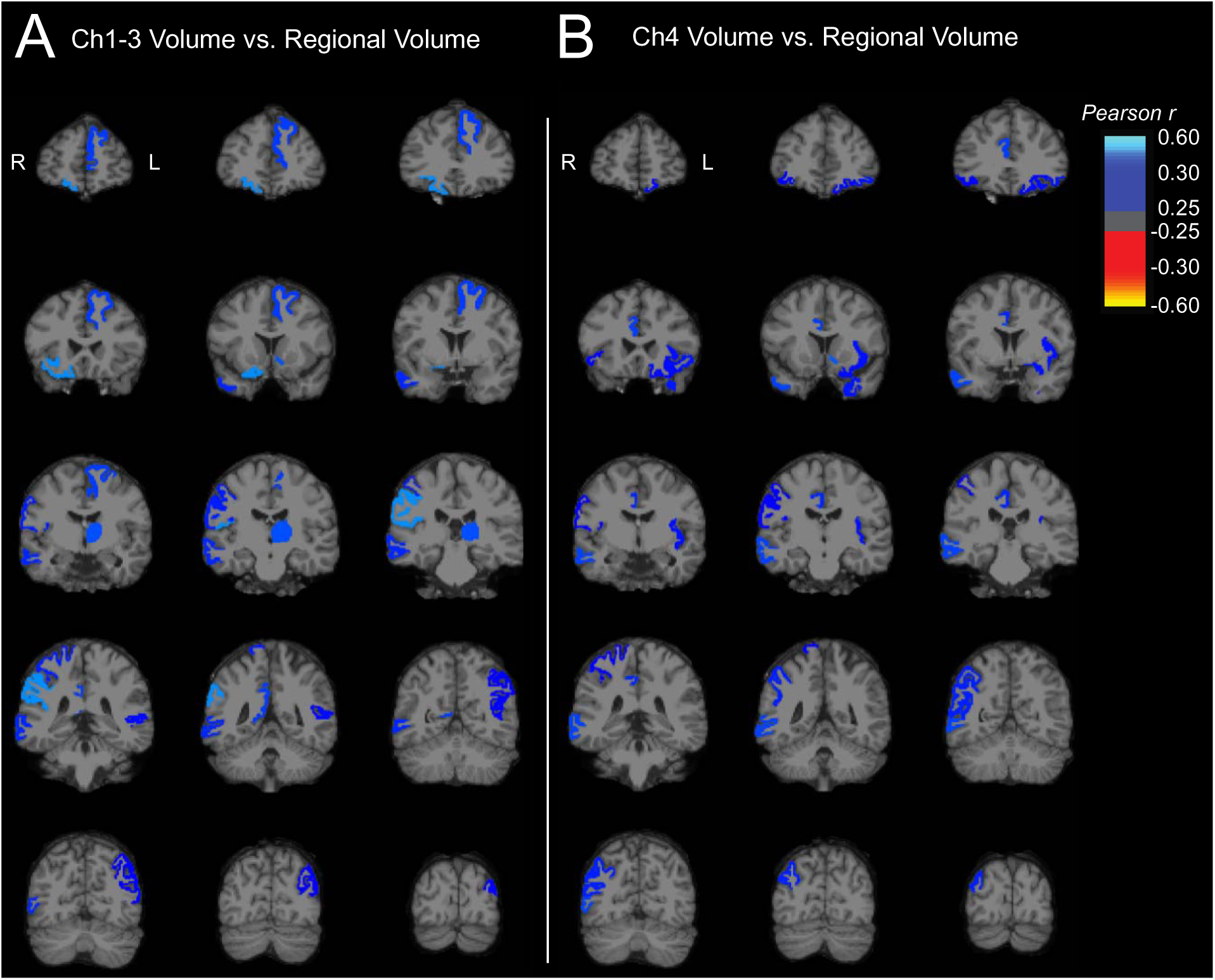
Brain maps of correlations between (A) Ch1-3 and (B) Ch4 basal forebrain volumes and FreeSurfer cortical and subcortical ROI volumes. Data depict Pearson r values for correlations between bilateral (A) Ch1-3 and (B) Ch4 volumes and the volume of each lateralized FreeSurfer [Version 6.0] ROI from the Desikan-Killiany atlas. Brain maps were created by producing images with the voxels in each FreeSurfer region set uniformly to the calculated Pearson r for that region and overlaid on an MNI template T1 MRI. Correlations were across all 84 lateralized FreeSurfer brain regions and displayed only for regions with uncorrected Pn<n0.05. MR image slices adhere to radiological convention, with orientation denoted in the first coronal section of each image series. Abbreviations: Ch1-3, cholinergic nuclei 1-3; Ch4, cholinergic nuclei.

## 4. DISCUSSION

This study aimed to further evaluate basal forebrain atrophy as a potential early marker of AD by examining the relationship between basal forebrain substructure volumes and cortical Aβ burden in middle-aged, cognitively normal older adults with varying *APOE*-indexed genetic risks of AD. Consistent with our *a priori* hypothesis and previous research^25,26^, we observed a significant inverse association between posterior basal forebrain volume (Ch4 or the nbM) and global cortical Aβ burden. Subgroup analyses that stratified by Aβ positivity (using *DVR* cutoffs of both 1.20 and 1.08) revealed this association was specific to Aβ+ individuals, with no significant relationship observed among those who were Aβ-. In contrast, anterior basal forebrain volumes were not significantly associated with cortical Aβ burden. Moreover, the volumes of medial-temporal regions, including hippocampus, parahippocampal gyrus, entorhinal cortex, and amygdala, showed no significant association with global cortical Aβ burden in this cognitively normal, at-risk cohort.

### 4.1. Basal forebrain atrophy as an early marker of Alzheimer’s disease

Acetylcholine is a neuro-transmitter critical for the modulation of learning, attention, and arousal.^46,47^ Impairments in acetylcholine synthesis and signaling – collectively referred to as cholinergic dysfunction – have been implicated in neurocognitive, behavioral, and psychological deficits observed in AD, forming the basis of the cholinergic hypothesis of AD.^48–51^ Subsequent research has suggested that cholinergic dysfunction in AD may be driven, at least in part, by the atrophy of the BFCS, the principal source of cortical acetylcholine in the central nervous system.^52,53^ Across the AD continuum, BFCS has been shown to undergo substantial neuronal loss, with reported volumetric reductions ranging from 8-87%.^17^ Notably, BFCS atrophy in AD appears to follow a spatially progressive trajectory, initiating in posterior regions – particularly the nucleus basalis of Meynert (Ch4) – and later advancing anteriorly to regions such as the medial septal nucleus and the diagonal band of Broca (Ch1-3).^17,26,54^ Among cognitively normal individuals with preclinical AD and those with MCI, atrophy tends to be largely confined to the nbM, sparing anterior substructures.^17,26^ In contrast, individuals with AD and related dementias typically exhibit more widespread BFCS atrophy, with the most pronounced volume loss occurring in the posterior Ch4 region.^17,54^ These findings have led to the proposal that posterior basal forebrain atrophy may serve as a sensitive early biomarker of AD pathogenesis.

Previous studies have shown that volume loss in the BFCS, particularly in the nbM, is associated with AD core pathologies among older adults without clinical symptoms^25,26^ and may even precede atrophy of the entorhinal cortex. ^22,23^ However, no prior study has examined the relationship between BFCS structural alterations and cortical Aβ load during the very early preclinical phases of AD in middle-aged adults, who may experience greater benefit from preventive strategies than older adults. Our study directly addresses this gap by demonstrating that nbM atrophy is associated with cortical Aβ burden in cognitively normal adults who are 50-67 years of age and Aβ+, but not in those who are Aβ-. This finding suggests that volume loss in the basal forebrain may occur as part of early AD pathogenesis, as indicated by amyloid plaque positivity. Furthermore, we found that *APOE* 1]4 homozygotes exhibited the highest levels of regional Aβ burden within the basal forebrain, consistent with the known increased susceptibility of ε4 carriers to Aβ plaque formation.^55^ Nonetheless, regional Aβ burden in basal forebrain substructures did not show significant inverse associations with their respective regional volumes, suggesting that the observed nbM atrophy is unlikely to be driven by local Aβ pathology, but may instead reflect the influence of existing AD pathogenesis.

### 4.2. Medial temporal atrophy cortical amyloid burden

The prevailing paradigm of AD centers on the accumulation of Aβ and tau pathologies,^56,57^ which are thought to drive neurodegeneration, typically first observed in the medial temporal lobe. However, emerging evidence suggests that basal forebrain atrophy may precede volume loss in the entorhinal cortex and hippocampus.^17,22,54,58^ In a previous analysis of this cohort, we found that global cortical Aβ burden was associated with global gray matter atrophy,^59^ but without characterization of which regions were driving the relationship. This paper further investigates the relationship between global cortical Aβ burden and the volumes of key medial temporal structures, including the hippocampus, parahippocampal gyrus, entorhinal cortex, and amygdala. In our cohort of middle-aged adults with varying levels of genetic risk for AD, cortical Aβ was not significantly associated with the volumes of the four medial temporal ROIs, irrespective of Aβ positivity. Further exploratory analyses examining all cortical ROIs found that middle temporal gyrus was the only cortical region whose volume was more strongly correlated with cortical Aβ burden than the nbM. Taken together, these findings provide added support for the hypothesis that nbM atrophy may serve as a robust, early biomarker of AD pathogenesis in the preclinical stage, potentially preceding degeneration of key medial temporal regions. Nevertheless, longitudinal studies are needed to empirically determine the temporal sequence of these regional structural changes.

### 4.3. Basal forebrain atrophy and projection areas

Mesulam and colleagues have delineated distinct projection patterns of basal forebrain substructures, showing that the anterior basal forebrain primarily innervates the hippocampus and olfactory bulb, while the posterior basal forebrain projects to the neocortex and amygdala.^6,7,45^ Given the central role of acetylcholine in neuromodulation,^27^ atrophy of basal forebrain substructures may lead to cholinergic deficits in their respective projection areas, potentially contributing to downstream neurodegeneration. However, in this study, we did not observe significant associations between lateralized volumes of the anterior basal forebrain and those of its primary projection target, the hippocampus. Similarly, lateralized posterior basal forebrain volumes were not broadly associated with neocortical or amygdala volumes, although we did find associations with a subset of frontal and temporal ROIs. One possible explanation for these null findings is the composition of our sample, which only included middle-aged, predominantly White, cognitively normal individuals without clinical diagnoses of MCI or dementia. As some of these individuals are in the preclinical stage of AD (i.e., Aβ+ with normal cognition), even though they may be experiencing cholinergic dysfunction and subtle neurocognitive changes, underlying pathologic changes may not yet be reflected in cortical atrophy detectable by structural MRI. Future studies should investigate whether basal forebrain atrophy affects the integrity of its projection areas using modalities more sensitive to early neuronal dysfunction, such as metabolic or synaptic PET imaging.

### 4.4. Limitations

This study is not without limitations. First, the tau status of participants was unknown, which limits our ability to contextualize basal forebrain atrophy within a more complete framework of AD pathology progression. ^20,21,60^ Second, concerns remain regarding the anatomical accuracy of basal forebrain localization, given the small size of its substructures and potential distortions introduced during image processing. However, the ROIs used in this study have been previously validated,^26,37^ and we took additional steps to ensure anatomical precision. Specifically, we visually inspected each basal forebrain ROI after transformation from MNI to participant-specific MR space to confirm correct anatomical placement and applied a SPM gray matter mask to all basal forebrain volumetric analyses. Although the modest sample size (n = 45) may limit the generalizability of our findings, participants were age- and sex-matched across *APOE* genotypes, reducing potential confounding and enhancing the validity of genotype-based comparisons. Finally, genetic risk was indexed by *APOE* genotype only. Future research should address these limitations through larger, longitudinal cohorts to clarify the temporal sequence of AD pathology in relation to basal forebrain and MTL degeneration across the AD continuum, incorporate tau PET imaging for more accurate disease staging, and adopt methods that allow for more detailed segmentation of basal forebrain substructures.^61^ Furthermore, while *APOE* genotype is by far the most important individual risk locus, genetic risk could also be studied by means of polygenic risk scores, which would incorporate more information.

### 4.5. Conclusions

In conclusion, this study of cognitively normal middle-aged adults at varying levels of genetic risk for AD demonstrated that posterior basal forebrain atrophy is associated with global cortical Aβ burden among Aβ+ individuals, but not among those who are Aβ-. In contrast, medial-temporal ROI volumes were not significantly associated with cortical Aβ burden, regardless of Aβ positivity, aligning with prior evidence suggesting that forebrain atrophy may precede medial-temporal volume loss during the course of AD progression. Taken together, these findings support the hypothesis that atrophy of the nucleus basalis of Meynert may serve as an early, preclinical biomarker of AD pathogenesis.

## Declarations

- Ethics approval and consent to participate: This research has been approved by the Institutional Review Board of Yale University and participants provided written informed consent.
- Consent for publication: Not applicable
- Availability of data and materials: All data used in this study can be provided upon reasonable request.
- Competing interests: Noted on the title page.
- Funding: Noted on the title page.
- Authors’ contributions: GC was involved in the analysis of data and writing of this manuscript. RO and AP were involved in data collection, data coordination, data analysis, and manuscript writing. KK was involved in data analysis and manuscript editing. CD and RC were involved in data collection and data coordination. BM was involved in manuscript writing.

## Acknowledgements

We thank the Yale ADRU staff for their contribution in participant recruitment and data collection.

