## Supplementary Files for "Basal forebrain volume is associated with cortical amyloid burden in cognitively unimpaired older adults at varying genetic risk for Alzheimer’s disease"

Yale University School of Medicine

One Church Street, 8^th^ Floor

New Haven, CT 06510

**Table of Contents:**

| **Content** | **Pages** |
| --- | --- |
| Supplementary Table 1. Participant characteristics stratified by Aβ positivity using a mean global cortical [^11^C]PiB *DVR* threshold of 1.2 | 3 |
| Supplementary Table 2. Participant characteristics stratified by Aβ positivity using a mean global cortical [^11^C]PiB *DVR* threshold of 1.08 | 4 |
| Supplementary Table 3. Relationship between basal forebrain substructural volumes and regional amyloid burden | 5 |
| Supplementary Table 4. Association between Freesurfer region volumes and global amyloid burden without partial volume correction | 6-7 |
| Supplementary Table 5. Association between Freesurfer region volumes and global amyloid burden with partial volume correction using Iterative Yang method | 8-9 |
| Supplementary Table 6. Relationship between basal forebrain and cortical volume across all FreeSurfer regions | 10-11 |
| Supplementary Figure 1. Mean global cortical amyloid burden by *APOE* genotype with (B) and without (A) partial volume correction | 12 |
| Supplementary Figure 2. Amyloid burden in basal forebrain substructures by *APOE* genotype after partial volume correction | 13 |
| Supplementary Figure 3. Association between partial volume-corrected mean global cortical amyloid burden and (A-C) Ch1-3 volume and (D-F) Ch4 volume | 14 |
| Supplementary Figure 4. Association between mean global cortical amyloid burden and (A-C) Ch1-3 volume and (D-F) Ch4 volume in Aβ + and Aβ- participants using a [^11^C]PiB *DVR* threshold of 1.2 | 15 |
| Supplementary Figure 5. Group differences in BFCS volume by amyloid status using a mean global cortical amyloid *DVR* threshold of 1.2 | 16 |
| Supplementary Figure 6. Association between mean global cortical amyloid burden and (A-C) Ch1-3 volume and (D-F) Ch4 volume in Aβ + and Aβ- participants using a [^11^C]PiB *DVR* threshold of 1.08 | 17 |
| Supplementary Figure 7. Association between partial volume-corrected mean global cortical amyloid burden and (A) hippocampal, (B) parahippocampal, (C) entorhinal, and (D) amygdala volume | 18 |
| Supplementary Figure 8. Association between mean global cortical amyloid burden and (A) hippocampal, (B) parahippocampal, (C) entorhinal, and (D) amygdalar volume in Aβ + and Aβ- participants using a [^11^C]PiB *DVR* threshold of 1.2 | 19 |
| Supplementary Figure 9. Group differences in medial temporal lobe volume by amyloid status using a mean global cortical amyloid *DVR* threshold of 1.2 | 20 |
| Supplementary Figure 10. Association between mean global cortical amyloid burden and (A) hippocampal, (B) parahippocampal, (C) entorhinal, and (D) amygdala volume in Aβ + and Aβ- participants using a [^11^C]PiB *DVR* threshold of 1.08 | 21 |

**Supplementary Table 1.** Participant characteristics stratified by Aβ positivity using a mean global cortical [^11^C]PiB *DVR* threshold of 1.2

|  | **Amyloid Negative** | **Amyloid Positive** | **t/χ^2^** | ***P*** |
| --- | --- | --- | --- | --- |
| **Participants (n)** | 34 | 11 |  |  |
| **Sex (M/F)** | 16/18 | 5/6 | 0.009 | 0.926 |
| **Age (years)** | 58.2 (4.6, 50.1-65.9) | 61.9 (4.1, 53.7-66.9) | 0.721 | 0.021* |
| **Education (years)** | 16.2 (1.8, 12.0-20.0) | 16.8 (2.0, 12.0-18.0) | 0.214 | 0.325 |
| **Global Amyloid (*DVR*)** | 1.03 (0.09, 0.91-1.20) | 1.39 (0.20, 1.20-1.77) | 7.316 | < 0.00001* |
| ***APOE* Genotype** |  |  | 6.738 | 0.034* |
| **ɛ3ɛ3** | 14 (41.2 %) | 1 (9.1 %) |  |  |
| **ɛ3ɛ4** | 12 (35.3 %) | 3 (27.3 %) |  |  |
| **ɛ4ɛ4** | 8 (23.5 %) | 7 (63.6 %) |  |  |

Data for continuous variables are mean (SD, range). *P* values are for unpaired t-tests (continuous variables) or *χ*^2^ tests (categorical variables). * *P* < 0.05. Abbreviations: *Aβ*, amyloid beta; *PiB*; Pittsburg compound B; *DVR*, distribution volume ratio; *APOE*, Apolipoprotein E.

**Supplementary Table 2.** Participant characteristics stratified by Aβ positivity using a mean global cortical [^11^C]PiB *DVR* threshold of 1.08

|  | **Amyloid Negative** | **Amyloid Positive** | **t/χ^2^** | ***P*** |
| --- | --- | --- | --- | --- |
| **Participants (n)** | 24 | 21 |  |  |
| **Sex (M/F)** | 9/15 | 12/9 | 1.74 | 0.19 |
| **Age (years)** | 57.45 (4.41, 50.06-64.84) | 61.02 (4.41, 51.03-66.86) | 7.33 | 0.01* |
| **Education (years)** | 16.08 (1.72, 12-18) | 16.62 (2.01 (12-20) | 0.93 | 0.34 |
| **Global Amyloid (*DVR*)** | 0.98 (0.05, 0.91-1.07) | 1.27 (0.19, 1.09-1.77) | 53.3 | < 0.0001* |
| **APOE ɛ4 Copy Number** |  |  | 13.39 | 0.001* |
| **0 copies** | 13 (54.2%) | 2 (9.5%) |  |  |
| **1 copy** | 8 (33.3%) | 7 (33.3%) |  |  |
| **2 copies** | 3 (12.5%) | 12 (57.1%) |  |  |

Data for continuous variables are mean (SD, range). *P* values are for unpaired t-tests (continuous variables) or *χ*^2^ tests (categorical variables). * *P* < 0.05. Abbreviations: *Aβ*, amyloid beta; *PiB*; Pittsburg compound B; *DVR*, distribution volume ratio; *APOE*, apolipoprotein E.

**Supplementary Table 3.** Relationship between basal forebrain substructural volumes and regional amyloid burden

| **Method of regional amyloid burden quantification** | **Basal forebrain substructures** | | | | | | | | | | | | | | | | |
| --- | --- | --- | --- | --- | --- | --- | --- | --- | --- | --- | --- | --- | --- | --- | --- | --- | --- |
|  | **Ch1-3, Left** | |  | **Ch1-3, Right** | |  | **Ch1-3, Bilateral** | |  | **Ch4, Left** | |  | **Ch4, Right** | |  | **Ch4, Bilateral** | |
|  | ***r*** | ***P*** |  | ***r*** | ***P*** |  | ***r*** | ***P*** |  | ***r*** | ***P*** |  | ***r*** | ***P*** |  | ***r*** | ***P*** |
| **Regional amyloid burden without PVC** | 0.21 | 0.16 |  | 0.30* | 0.05 |  | 0.29 | 0.05 |  | 0.05 | 0.77 |  | 0.003 | 0.98 |  | 0.01 | 0.96 |
| **Regional amyloid burden after IY-PVC** | 0.20 | 0.18 |  | 0.30* | 0.05 |  | 0.30* | 0.05 |  | -0.01 | 0.94 |  | -0.10 | 0.51 |  | -0.12 | 0.42 |

Acronyms. *Ch*, Cholinergic nuclei; *IY-PVC*, Iterative Yang partial volume correction; r = Pearson r

**Supplementary Table 4.** Association between Freesurfer region volumes and global amyloid burden without partial volume correction

| **Bilateral region** | **Full sample** | | **Amyloid positive** | | **Amyloid negative** | |
| --- | --- | --- | --- | --- | --- | --- |
|  | **Pearson *r*** | ***P*** | **Pearson *r*** | ***P*** | **Pearson *r*** | ***P*** |
| bl_frontalpole | -0.29 | 0.06 | -0.09 | 0.78 | -0.29 | 0.09 |
| bl_superiorfrontal | -0.21 | 0.18 | 0.30 | 0.37 | 0.16 | 0.37 |
| bl_rostralmiddlefrontal | -0.21 | 0.17 | -0.11 | 0.74 | 0.02 | 0.93 |
| bl_caudalmiddlefrontal | -0.13 | 0.39 | 0.08 | 0.81 | 0.09 | 0.63 |
| bl_parsorbitalis | -0.26 | 0.08 | -0.36 | 0.28 | -0.20 | 0.27 |
| bl_parsopercularis | -0.12 | 0.43 | 0.35 | 0.29 | 0.13 | 0.45 |
| bl_parstriangularis | 0.10 | 0.53 | 0.12 | 0.73 | 0.16 | 0.35 |
| bl_lateralorbitofrontal | -0.35* | 0.02 | -0.49 | 0.12 | -0.28 | 0.11 |
| bl_medialorbitofrontal | -0.27 | 0.08 | -0.28 | 0.41 | -0.35* | 0.05 |
| bl_temporalpole | -0.35* | 0.02 | -0.57 | 0.07 | -0.13 | 0.46 |
| bl_entorhinal | -0.05 | 0.73 | -0.49 | 0.13 | -0.03 | 0.85 |
| bl_parahippocampal | -0.26 | 0.08 | -0.31 | 0.36 | -0.11 | 0.54 |
| bl_Hippocampus | -0.15 | 0.33 | 0.58 | 0.06 | -0.15 | 0.40 |
| bl_Amygdala | -0.02 | 0.91 | 0.31 | 0.35 | -0.13 | 0.47 |
| bl_inferiortemporal | 0.25 | 0.10 | 0.41 | 0.21 | -0.02 | 0.91 |
| bl_fusiform | -0.18 | 0.23 | -0.43 | 0.19 | -0.03 | 0.88 |
| bl_middletemporal | -0.49* | 0.0006 | -0.36 | 0.28 | -0.24 | 0.18 |
| bl_bankssts | -0.32* | 0.03 | 0.06 | 0.87 | -0.19 | 0.28 |
| bl_superiortemporal | -0.34* | 0.02 | -0.33 | 0.32 | -0.23 | 0.18 |
| bl_transversetemporal | 0.09 | 0.55 | 0.27 | 0.43 | -0.04 | 0.82 |
| bl_supramarginal | -0.07 | 0.66 | 0.12 | 0.73 | 0.10 | 0.56 |
| bl_insula | -0.33* | 0.03 | -0.55 | 0.08 | -0.10 | 0.57 |
| bl_rostralanteriorcingulate | -0.35* | 0.02 | -0.53 | 0.09 | 0.00 | 1.00 |
| bl_caudalanteriorcingulate | -0.33* | 0.03 | -0.31 | 0.36 | -0.42* | 0.01 |
| bl_posteriorcingulate | -0.32* | 0.03 | -0.34 | 0.31 | -0.02 | 0.91 |
| bl_isthmuscingulate | -0.27 | 0.07 | -0.01 | 0.99 | -0.08 | 0.63 |
| bl_precuneus | -0.03 | 0.84 | 0.04 | 0.90 | 0.01 | 0.96 |
| bl_paracentral | -0.26 | 0.09 | 0.20 | 0.56 | -0.18 | 0.31 |
| bl_postcentral | 0.06 | 0.68 | 0.81* | 0.003 | 0.14 | 0.43 |
| bl_precentral | -0.02 | 0.90 | 0.28 | 0.41 | -0.10 | 0.58 |
| bl_superiorparietal | 0.07 | 0.66 | 0.11 | 0.75 | 0.04 | 0.80 |
| bl_inferiorparietal | -0.28 | 0.07 | 0.33 | 0.33 | -0.35* | 0.04 |
| bl_lateraloccipital | -0.27 | 0.07 | -0.09 | 0.80 | -0.18 | 0.31 |
| bl_cuneus | -0.28 | 0.06 | -0.41 | 0.22 | -0.22 | 0.21 |
| bl_pericalcarine | -0.13 | 0.41 | -0.08 | 0.81 | -0.18 | 0.30 |
| bl_lingual | -0.29 | 0.05 | -0.19 | 0.58 | -0.19 | 0.28 |
| bl_Thalamus_Proper | -0.35* | 0.02 | 0.08 | 0.82 | -0.44 | 0.01 |
| bl_Caudate | -0.05 | 0.76 | 0.10 | 0.77 | -0.10 | 0.59 |
| bl_Putamen | -0.24 | 0.12 | 0.12 | 0.73 | -0.32 | 0.07 |
| bl_Pallidum | -0.20 | 0.18 | 0.05 | 0.89 | -0.45* | 0.01 |
| bl_Accumbens_area | -0.24 | 0.12 | -0.01 | 0.97 | -0.18 | 0.30 |
| bl_VentralDC | -0.24 | 0.12 | -0.11 | 0.76 | -0.42* | 0.01 |

**Supplementary Table 5.** Association between Freesurfer region volumes and global amyloid burden with partial volume correction using Iterative Yang method

| **Bilateral region** | **Full sample** | | **Amyloid positive** | | **Amyloid negative** | |
| --- | --- | --- | --- | --- | --- | --- |
|  | **Pearson *r*** | ***P*** | **Pearson *r*** | ***P*** | **Pearson *r*** | ***P*** |
| bl_frontalpole | -0.34* | 0.02 | -0.18 | 0.40 | -0.41 | 0.07 |
| bl_superiorfrontal | -0.26 | 0.08 | -0.35 | 0.09 | -0.47* | 0.03 |
| bl_rostralmiddlefrontal | -0.23 | 0.13 | -0.33 | 0.11 | -0.50* | 0.02 |
| bl_caudalmiddlefrontal | -0.20 | 0.18 | -0.32 | 0.12 | -0.16 | 0.48 |
| bl_parsorbitalis | -0.26 | 0.09 | -0.16 | 0.46 | -0.26 | 0.25 |
| bl_parsopercularis | -0.12 | 0.42 | -0.16 | 0.46 | -0.15 | 0.51 |
| bl_parstriangularis | 0.10 | 0.53 | -0.01 | 0.98 | -0.32 | 0.16 |
| bl_lateralorbitofrontal | -0.32* | 0.03 | -0.20 | 0.34 | -0.52* | 0.02 |
| bl_medialorbitofrontal | -0.29* | 0.05 | -0.12 | 0.59 | -0.21 | 0.35 |
| bl_temporalpole | -0.34* | 0.02 | -0.32 | 0.13 | -0.20 | 0.38 |
| bl_entorhinal | -0.03 | 0.85 | -0.26 | 0.23 | -0.32 | 0.15 |
| bl_parahippocampal | -0.23 | 0.13 | -0.15 | 0.50 | -0.22 | 0.33 |
| bl_Hippocampus | -0.16 | 0.30 | 0.10 | 0.65 | -0.32 | 0.15 |
| bl_Amygdala | 0.02 | 0.91 | 0.15 | 0.49 | -0.33 | 0.14 |
| bl_inferiortemporal | 0.24 | 0.12 | 0.21 | 0.31 | -0.18 | 0.43 |
| bl_fusiform | -0.21 | 0.17 | -0.17 | 0.43 | -0.05 | 0.83 |
| bl_middletemporal | -0.49* | 0.0006 | -0.49* | 0.02 | -0.13 | 0.59 |
| bl_bankssts | -0.35* | 0.02 | -0.27 | 0.20 | -0.18 | 0.43 |
| bl_superiortemporal | -0.32* | 0.03 | -0.27 | 0.21 | -0.35 | 0.12 |
| bl_transversetemporal | 0.13 | 0.40 | 0.23 | 0.28 | -0.25 | 0.27 |
| bl_supramarginal | -0.09 | 0.55 | -0.17 | 0.41 | -0.35 | 0.12 |
| bl_insula | -0.35* | 0.02 | -0.55* | 0.005 | -0.04 | 0.87 |
| bl_rostralanteriorcingulate | -0.37* | 0.01 | -0.52 | 0.01 | -0.24 | 0.29 |
| bl_caudalanteriorcingulate | -0.34* | 0.02 | -0.17 | 0.42 | -0.46* | 0.03 |
| bl_posteriorcingulate | -0.32* | 0.03 | -0.28 | 0.18 | -0.19 | 0.40 |
| bl_isthmuscingulate | -0.27 | 0.08 | -0.25 | 0.25 | 0.02 | 0.92 |
| bl_precuneus | -0.07 | 0.63 | -0.16 | 0.46 | -0.15 | 0.53 |
| bl_paracentral | -0.27 | 0.07 | -0.10 | 0.64 | -0.41 | 0.07 |
| bl_postcentral | 0.03 | 0.84 | 0.13 | 0.53 | -0.08 | 0.72 |
| bl_precentral | -0.07 | 0.64 | 0.03 | 0.90 | 0.01 | 0.96 |
| bl_superiorparietal | 0.04 | 0.77 | 0.06 | 0.79 | -0.17 | 0.45 |
| bl_inferiorparietal | -0.29 | 0.05 | -0.10 | 0.65 | -0.13 | 0.58 |
| bl_lateraloccipital | -0.25 | 0.09 | -0.15 | 0.49 | -0.13 | 0.58 |
| bl_cuneus | -0.30* | 0.04 | -0.31 | 0.14 | 0.15 | 0.52 |
| bl_pericalcarine | -0.15 | 0.34 | -0.07 | 0.76 | 0.25 | 0.27 |
| bl_lingual | -0.26 | 0.08 | -0.16 | 0.45 | 0.42 | 0.06 |
| bl_Thalamus_Proper | -0.37* | 0.01 | -0.23 | 0.27 | -0.58 | 0.01 |
| bl_Caudate | 0.00 | 0.98 | 0.21 | 0.31 | -0.04 | 0.86 |
| bl_Putamen | -0.18 | 0.23 | 0.08 | 0.70 | -0.21 | 0.37 |
| bl_Pallidum | -0.19 | 0.20 | 0.13 | 0.54 | -0.09 | 0.68 |
| bl_Accumbens_area | -0.20 | 0.18 | -0.08 | 0.71 | -0.37 | 0.10 |
| bl_VentralDC | -0.22 | 0.15 | 0.05 | 0.82 | -0.24 | 0.30 |

| **Supplementary Table 6.** Relationship between basal forebrain and cortical volume across all FreeSurfer regions | | | | | |
| --- | --- | --- | --- | --- | --- |
| **Left Hemisphere Region** | **Ch1-3**  **Pearson *r* (*P*)** | **Ch4**  **Pearson *r* (*P*)** | **Right Hemisphere Region** | **Ch1-3**  **Pearson *r* (*P*)** | **Ch4**  **Pearson *r* (*P*)** |
| ctx-lh-frontalpole | -0.08 (0.59) | 0.32 (0.03*) | ctx-rh-frontalpole | 0.16 (0.29) | 0.19 (0.21) |
| ctx-lh-superiorfrontal | 0.37 (0.01*) | 0.16 (0.28) | ctx-rh-superiorfrontal | 0.25 (0.10) | 0.13 (0.40) |
| ctx-lh-rostralmiddlefrontal | 0.26 (0.09) | 0.14 (0.36) | ctx-rh-rostralmiddlefrontal | 0.24 (0.11) | 0.11 (0.49) |
| ctx-lh-caudalmiddlefrontal | 0.21 (0.17) | 0.06 (0.69) | ctx-rh-caudalmiddlefrontal | 0.28 (0.07) | 0.04 (0.78) |
| ctx-lh-parsorbitalis | -0.01 (0.98) | 0.33 (0.03*) | ctx-rh-parsorbitalis | 0.06 (0.69) | 0.30 (0.05*) |
| ctx-lh-parsopercularis | 0.04 (0.79) | -0.16 (0.30) | ctx-rh-parsopercularis | 0.04 (0.81) | 0.05 (0.72) |
| ctx-lh-parstriangularis | 0.15 (0.32) | -0.16 (0.29) | ctx-rh-parstriangularis | -0.15 (0.33) | -0.16 (0.29) |
| ctx-lh-lateralorbitofrontal | 0.22 (0.14) | 0.31 (0.04*) | ctx-rh-lateralorbitofrontal | 0.46 (0.001*) | 0.27 (0.07) |
| ctx-lh-medialorbitofrontal | -0.22 (0.15) | 0.11 (0.48) | ctx-rh-medialorbitofrontal | 0.18 (0.25) | 0.06 (0.70) |
| ctx-lh-temporalpole | -0.22 (0.15) | 0.33 (0.03*) | ctx-rh-temporalpole | -0.10 (0.52) | 0.21 (0.16) |
| ctx-lh-entorhinal | -0.09 (0.54) | 0.08 (0.60) | ctx-rh-entorhinal | 0.05 (0.74) | 0.27 (0.07) |
| ctx-lh-parahippocampal | 0.15 (0.34) | 0.19 (0.21) | ctx-rh-parahippocampal | 0.27 (0.08) | 0.28 (0.06) |
| Left-Hippocampus | 0.19 (0.21) | 0.22 (0.15) | Right-Hippocampus | 0.04 (0.79) | 0.17 (0.25) |
| Left-Amygdala | 0.23 (0.12) | 0.04 (0.78) | Right-Amygdala | 0.11 (0.46) | -0.05 (0.75) |
| ctx-lh-inferiortemporal | -0.19 (0.20) | -0.06 (0.69) | ctx-rh-inferiortemporal | -0.06 (0.71) | -0.11 (0.46) |
| ctx-lh-fusiform | -0.16 (0.29) | -0.02 (0.88) | ctx-rh-fusiform | -0.19 (0.20) | -0.06 (0.72) |
| ctx-lh-middletemporal | 0.09 (0.54) | 0.11 (0.46) | ctx-rh-middletemporal | 0.35 (0.02*) | 0.38 (0.01*) |
| ctx-lh-bankssts | 0.31 (0.04*) | 0.20 (0.19) | ctx-rh-bankssts | 0.21 (0.18) | 0.07 (0.65) |
| ctx-lh-superiortemporal | 0.18 (0.25) | 0.14 (0.37) | ctx-rh-superiortemporal | 0.10 (0.52) | -0.03 (0.86) |
| ctx-lh-transversetemporal | 0.12 (0.42) | -0.01 (0.97) | ctx-rh-transversetemporal | -0.05 (0.74) | -0.27 (0.08) |
| ctx-lh-supramarginal | 0.24 (0.11) | 0.19 (0.21) | ctx-rh-supramarginal | 0.47 (0.001*) | -0.07 (0.63) |
| ctx-lh-insula | 0.27 (0.08) | 0.32 (0.03*) | ctx-rh-insula | -0.04 (0.81) | 0.14 (0.35) |
| ctx-lh-rostralanteriorcingulate | 0.07 (0.64) | 0.09 (0.56) | ctx-rh-rostralanteriorcingulate | 0.10 (0.52) | 0.26 (0.08) |
| ctx-lh-caudalanteriorcingulate | -0.01 (0.96) | 0.10 (0.50) | ctx-rh-caudalanteriorcingulate | 0.09 (0.58) | 0.37 (0.01*) |
| ctx-lh-posteriorcingulate | 0.04 (0.79) | 0.19 (0.20) | ctx-rh-posteriorcingulate | 0.07 (0.63) | 0.35 (0.02*) |
| ctx-lh-isthmuscingulate | 0.21 (0.17) | 0.07 (0.67) | ctx-rh-isthmuscingulate | 0.37 (0.01*) | 0.17 (0.26) |
| ctx-lh-precuneus | 0.19 (0.23) | -0.03 (0.82) | ctx-rh-precuneus | 0.17 (0.25) | 0.06 (0.70) |
| ctx-lh-paracentral | 0.18 (0.23) | 0.22 (0.14) | ctx-rh-paracentral | 0.06 (0.69) | -0.05 (0.74) |
| ctx-lh-postcentral | 0.15 (0.33) | 0.13 (0.39) | ctx-rh-postcentral | 0.33 (0.03*) | 0.30 (0.05*) |
| ctx-lh-precentral | 0.27 (0.07) | 0.22 (0.15) | ctx-rh-precentral | 0.12 (0.44) | 0.17 (0.26) |
| ctx-lh-superiorparietal | 0.24 (0.12) | -0.13 (0.39) | ctx-rh-superiorparietal | 0.19 (0.21) | -0.24 (0.11) |
| ctx-lh-inferiorparietal | 0.30 (0.05*) | 0.16 (0.30) | ctx-rh-inferiorparietal | 0.12 (0.45) | 0.34 (0.02*) |
| ctx-lh-lateraloccipital | -0.16 (0.31) | 0.09 (0.56) | ctx-rh-lateraloccipital | -0.13 (0.41) | 0.25 (0.10) |
| ctx-lh-cuneus | 0.01 (0.95) | 0.07 (0.67) | ctx-rh-cuneus | -0.08 (0.58) | -0.07 (0.66) |
| ctx-lh-pericalcarine | -0.01 (0.94) | -0.01 (0.94) | ctx-rh-pericalcarine | 0.01 (0.95) | 0.01 (0.94) |
| ctx-lh-lingual | 0.22 (0.14) | 0.11 (0.47) | ctx-rh-lingual | 0.07 (0.66) | 0.08 (0.62) |
| Left-Thalamus-Proper | 0.39 (0.01*) | 0.23 (0.13) | Right-Thalamus-Proper | 0.19 (0.23) | 0.24 (0.11) |
| Left-Caudate | 0.09 (0.58) | 0.29 (0.05*) | Right-Caudate | 0.11 (0.49) | 0.23 (0.12) |
| Left-Putamen | 0.11 (0.48) | 0.07 (0.64) | Right-Putamen | 0.22 (0.14) | 0.24 (0.11) |
| Left-Pallidum | 0.22 (0.16) | 0.29 (0.05*) | Right-Pallidum | 0.19 (0.22) | 0.25 (0.09) |
| Left-Accumbens-area | 0.40 (0.01*) | 0.40 (0.01*) | Right-Accumbens-area | 0.21 (0.16) | 0.15 (0.34) |
| Left-VentralDC | 0.06 (0.70) | 0.09 (0.56) | Right-VentralDC | 0.00 (0.98) | 0.02 (0.91) |

Data are Pearson *r* (*P* values) for correlations between bilateral Ch1-3 and Ch4 volumes and the volume of each lateralized FreeSurfer [Version 6.0] ROI from the Desikan-Killiany atlas. * *P* < 0.05. Abbreviations: *Ch1-3*, cholinergic nuclei 1-3; *Ch4*, cholinergic nuclei.

**Supplementary Figure 1.**

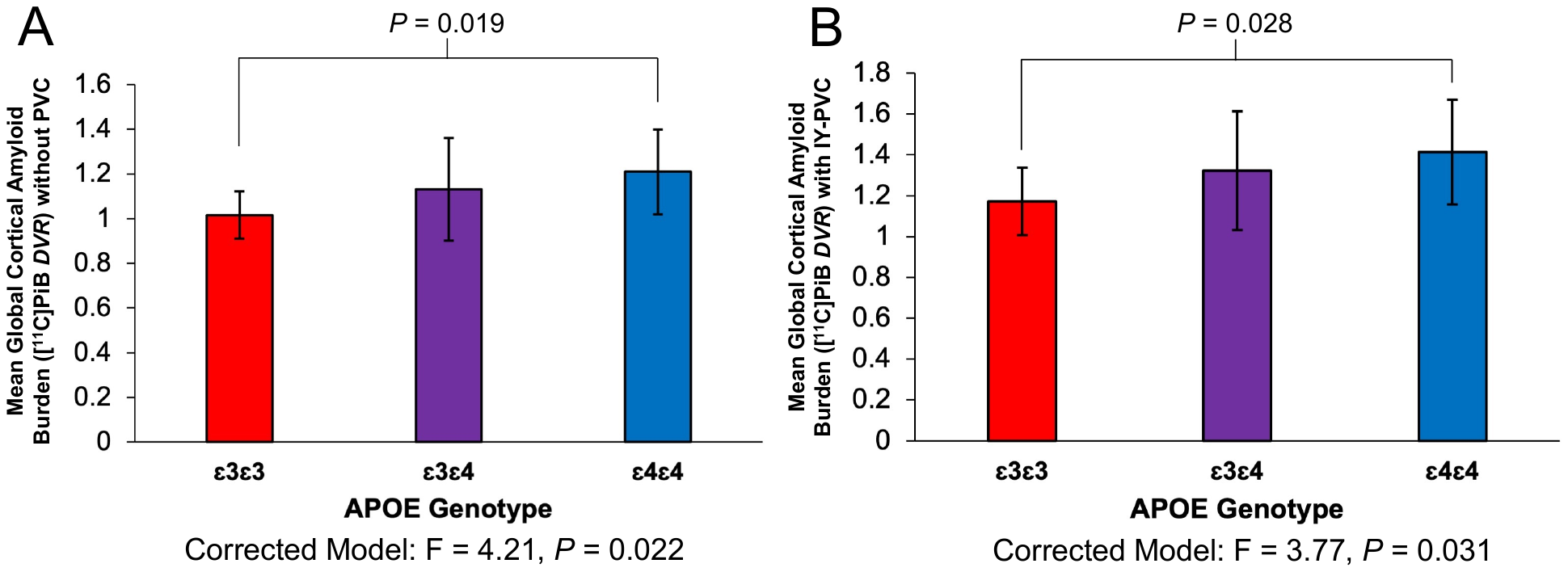

Mean global cortical amyloid burden by *APOE* genotype with (B) and without (A) partial volume correction. Group differences in global cortical amyloid burden by *APOE* genotype were calculated using one-way ANOVA with post-hoc Bonferroni unpaired t-tests. *F* statistics and *P* values for each model are depicted below the x-axis and *P* values from post-hoc unpaired t-tests are depicted above the bar graphs to denote significant group differences. Global cortical Aβ burden was calculated as a weighted average of composite regions commonly affected by Aβ burden in AD, which included prefrontal, lateral temporal, posterior cingulate/precuneus, and lateral parietal ROIs. Abbreviations: *Aβ*, amyloid beta; *AD*, Alzheimer’s disease; *APOE*, apolipoprotein E; *PiB*; Pittsburg compound B; *IY-PVC*, Iterative Yang partial volume correction; *DVR*, distribution volume ratio.

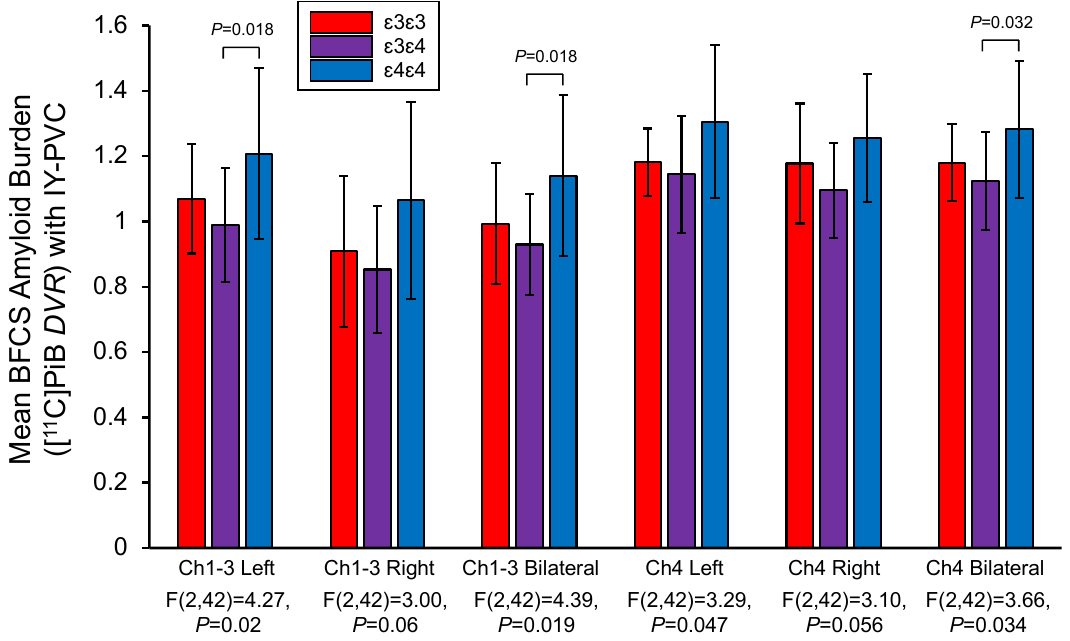
**Supplementary Figure 2.**

Amyloid burden in basal forebrain substructures by *APOE* genotype after partial volume correction. Group differences in BFCS amyloid burden by *APOE* genotype were calculated using one-way ANOVA with post-hoc Bonferroni unpaired t-tests. *F* statistics and *P* values for each model are depicted below the x-axis and *P* values from post-hoc unpaired t-tests are depicted above the bar graphs to denote significant group differences. Abbreviations: *BFCS*, basal forebrain cholinergic system; *PiB*; Pittsburg compound B; *IY-PVC*, Iterative Yang partial volume correction; *DVR*, distribution volume ratio; *Ch1-3*, cholinergic nuclei 1-3; *Ch4*, cholinergic nuclei 4.

**Supplementary Figure 3.**

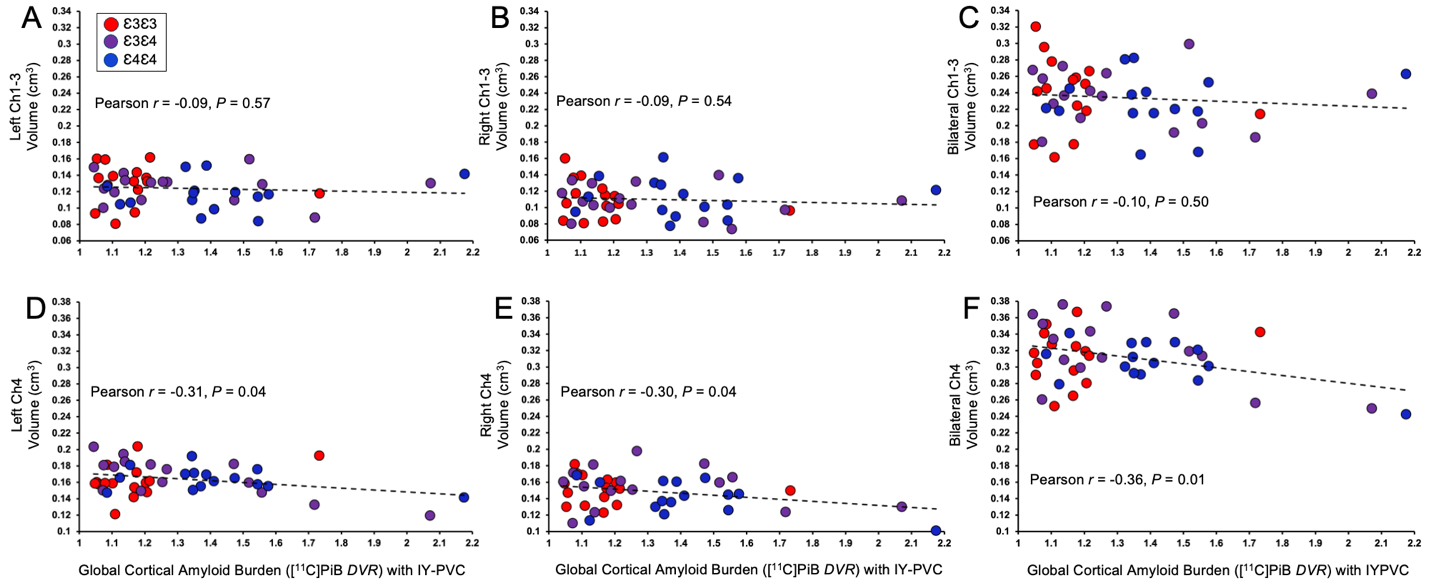

Association between partial volume-corrected mean global cortical amyloid burden and (A-C) Ch1-3 volume and (D-F) Ch4 volume. Displayed are Pearson *r* (*P*) for correlations between global cortical amyloid burden *(DVR)* and basal forebrain volumes. Global cortical Aβ burden was calculated as a weighted average of composite regions commonly affected by Aβ burden in AD, which included prefrontal, lateral temporal, posterior cingulate/precuneus, and lateral parietal ROIs. Abbreviations: *Aβ*, amyloid beta; *AD*, Alzheimer’s disease; *IY-PVC*, Iterative Yang partial volume correction; *PiB*; Pittsburg compound B; *DVR*, distribution volume ratio; *Ch1-3*, cholinergic nuclei 1-3; *Ch4*, cholinergic nuclei 4.

**Supplementary Figure 4.**

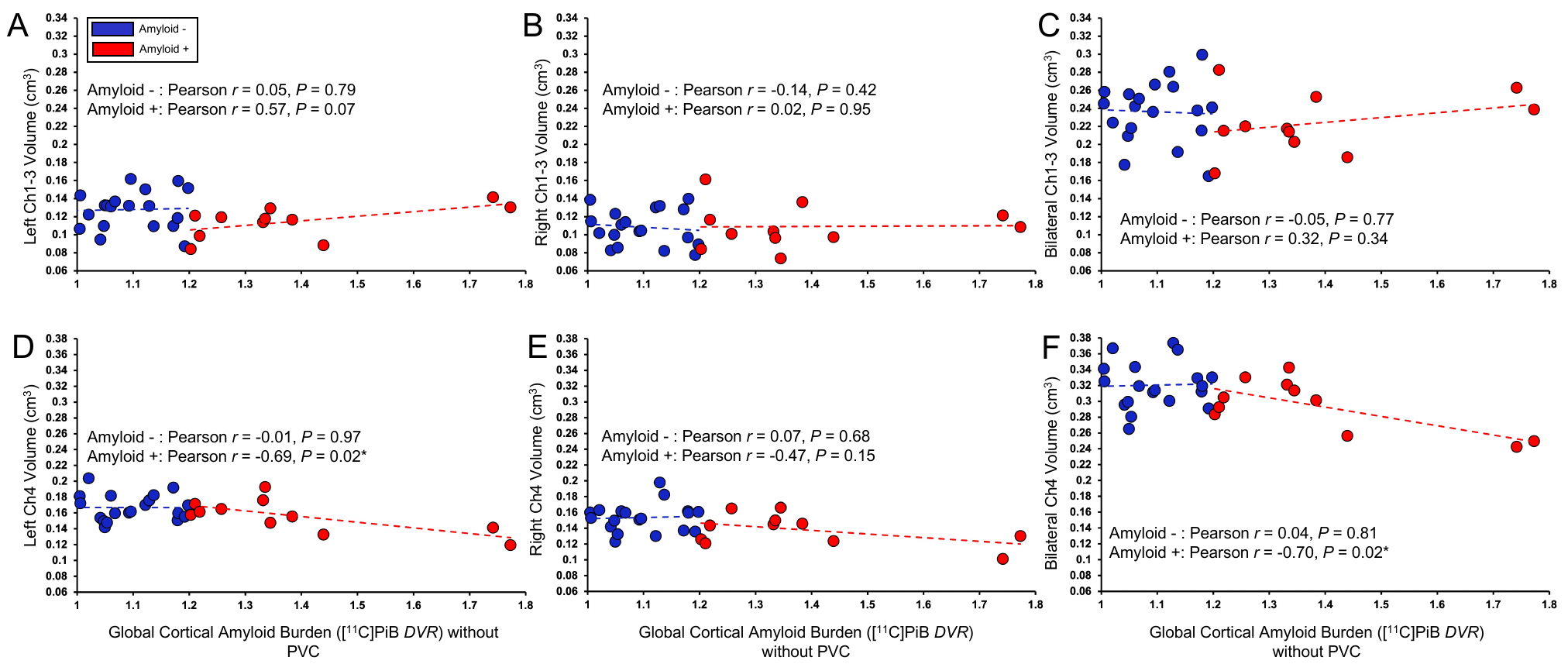

Association between mean global cortical amyloid burden and (A-C) Ch1-3 volume and (D-F) Ch4 volume in Aβ + and Aβ- participants using a [^11^C]PiB *DVR* threshold of 1.2. Data are Pearson *r* (*P*) for correlations between mean global cortical amyloid burden (*DVR*) and basal forebrain volumes, with separate models for Aβ + and Aβ- participants. A *DVR* threshold of 1.2 was used to determine amyloid status (positive vs. negative). Global cortical Aβ burden was calculated as a weighted average of composite regions commonly affected by Aβ burden in AD, which included prefrontal, lateral temporal, posterior cingulate/precuneus, and lateral parietal ROIs. Abbreviations: *Aβ*, amyloid beta; *AD*, Alzheimer’s disease; *PiB*; Pittsburg compound B; *DVR*, distribution volume ratio; *Ch1-3*, cholinergic nuclei 1-3; *Ch4*, cholinergic nuclei 4.

**Supplementary Figure 5.**

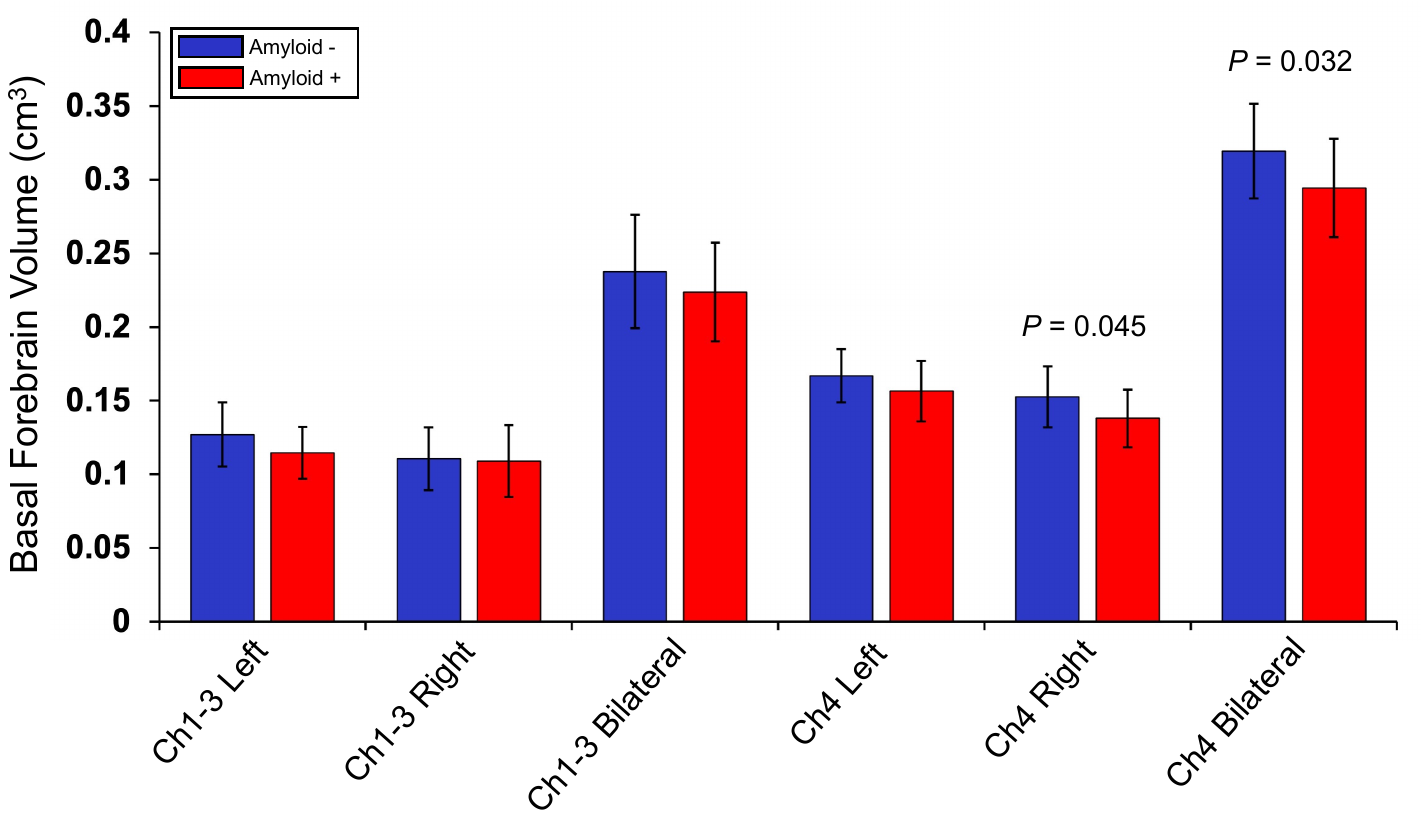

Group differences in BFCS volume by amyloid status using a mean global cortical amyloid *DVR* threshold of 1.2. Bar graphs represent mean (SD) for Ch1-3 and Ch4 basal forebrain volumes (cm^3^) for amyloid positive and negative participants. Volumes were normalized using each participant’s eICV. Group differences were determined using unpaired t-tests and *P* values are depicted above the bar graphs to denote significant group differences. A mean global cortical [^11^C]PiB *DVR* threshold of 1.2 was used to determine amyloid status (positive vs. negative). Global cortical Aβ burden was calculated as a weighted average of composite regions commonly affected by Aβ burden in AD, which included prefrontal, lateral temporal, posterior cingulate/precuneus, and lateral parietal ROIs. Abbreviations: *Aβ*, amyloid beta; *AD*, Alzheimer’s disease; *PiB*; Pittsburg compound B; *DVR*, distribution volume ratio; *eICV*, estimated intracranial volume; *Ch1-3*, cholinergic nuclei 1-3; *Ch4*, cholinergic nuclei 4.

**Supplementary Figure 6.**

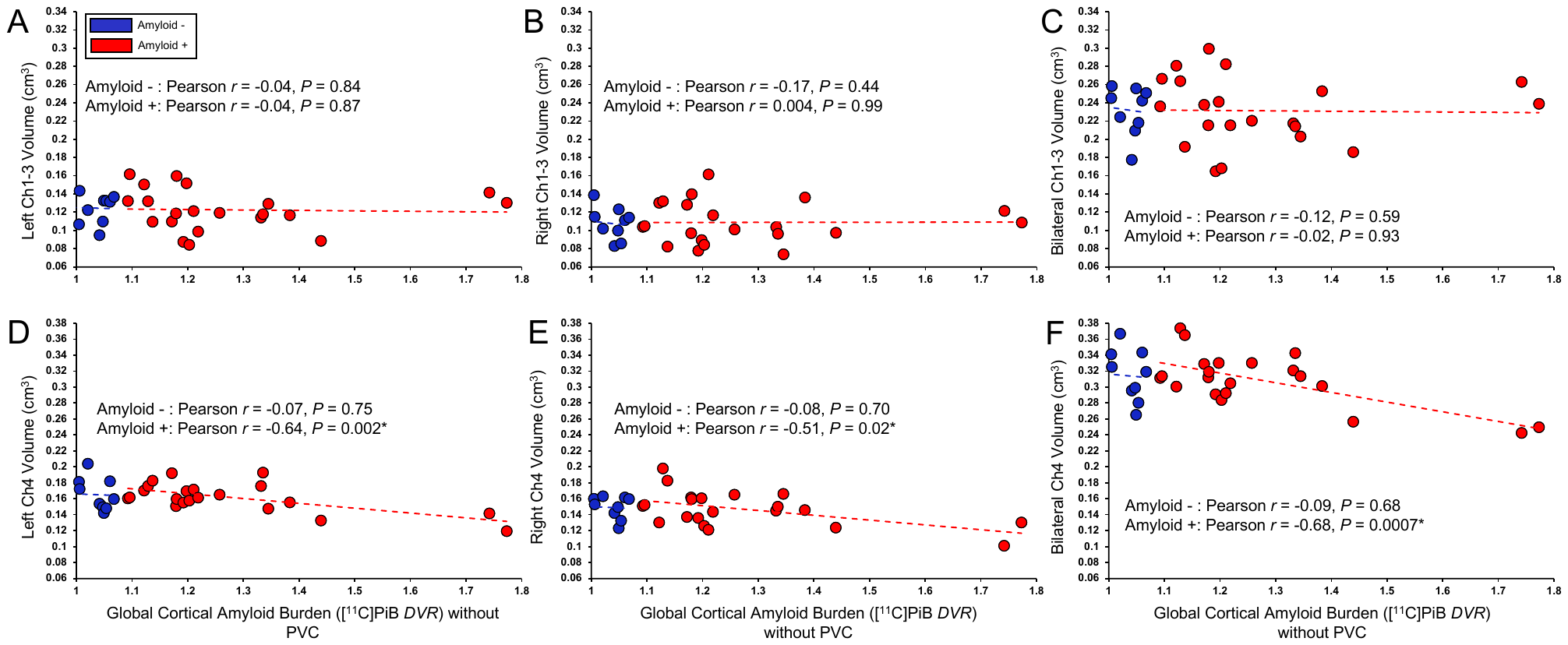

Association between mean global cortical amyloid burden and (A-C) Ch1-3 volume and (D-F) Ch4 volume in Aβ + and Aβ- participants using a [^11^C]PiB *DVR* threshold of 1.08. Data are Pearson *r* (*P*) for correlations between mean global cortical amyloid burden (*DVR*) and basal forebrain volumes, with separate models for Aβ + and Aβ- participants. A *DVR* threshold of 1.08 was used to determine amyloid status (positive vs. negative). Global cortical Aβ burden was calculated as a weighted average of composite regions commonly affected by Aβ burden in AD, which included prefrontal, lateral temporal, posterior cingulate/precuneus, and lateral parietal ROIs. Abbreviations: *Aβ*, amyloid beta; *AD*, Alzheimer’s disease; *PiB*; Pittsburg compound B; *DVR*, distribution volume ratio; *Ch1-3*, cholinergic nuclei 1-3; *Ch4*, cholinergic nuclei 4.

**Supplementary Figure 7.**

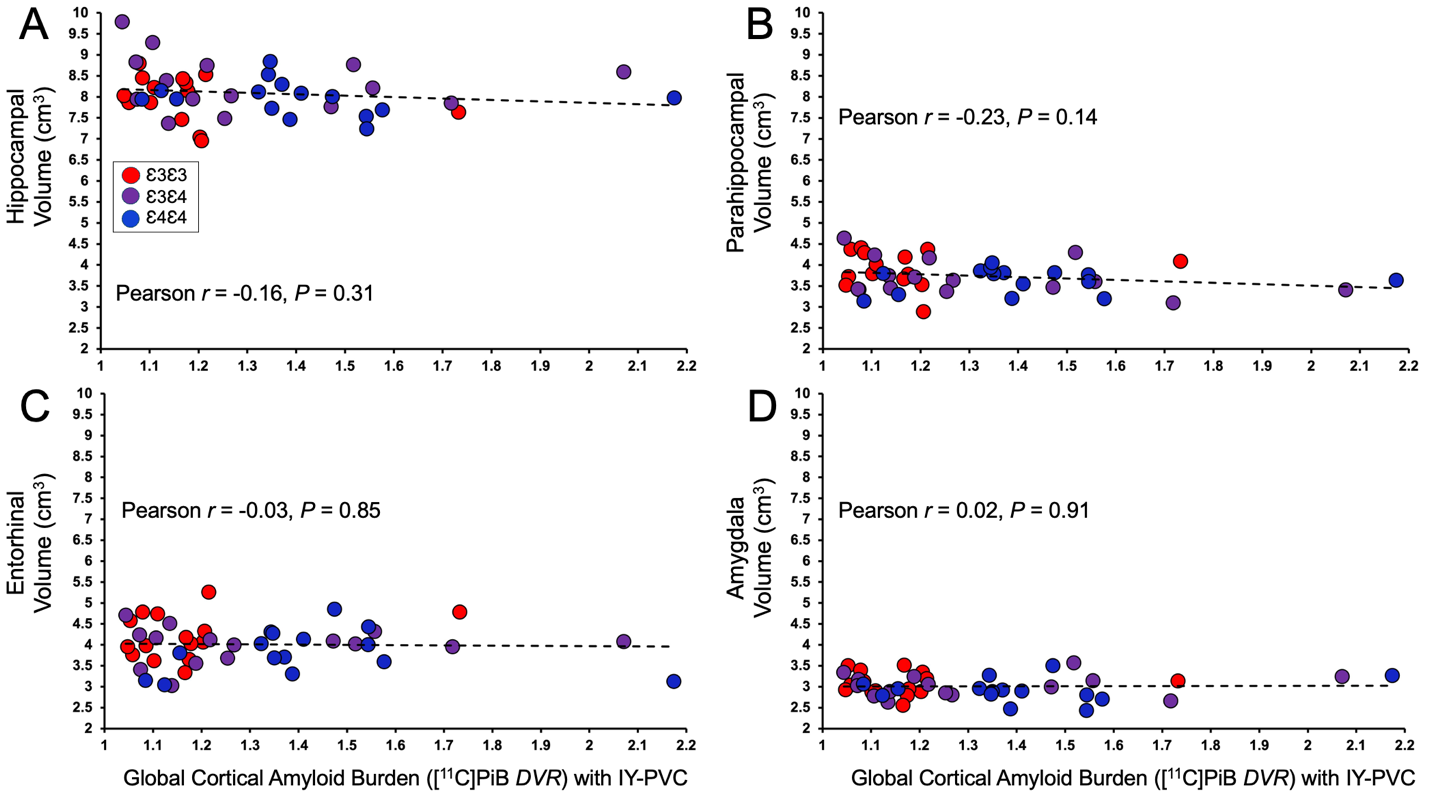

Association between partial volume-corrected mean global cortical amyloid burden and (A) hippocampal, (B) parahippocampal, (C) entorhinal, and (D) amygdalar volume. Displayed are Pearson *r* (*P*) for correlations between global cortical amyloid burden *(DVR)* and medial temporal lobe volumes. Global cortical Aβ burden was calculated as a weighted average of composite regions commonly affected by Aβ burden in AD, which included prefrontal, lateral temporal, posterior cingulate/precuneus, and lateral parietal ROIs. Abbreviations: *Aβ*, amyloid beta; *AD*, Alzheimer’s disease; *IY-PVC*, Iterative Yang partial volume correction; *PiB*; Pittsburg compound B; *DVR*, distribution volume ratio.

**Supplementary Figure 8.**

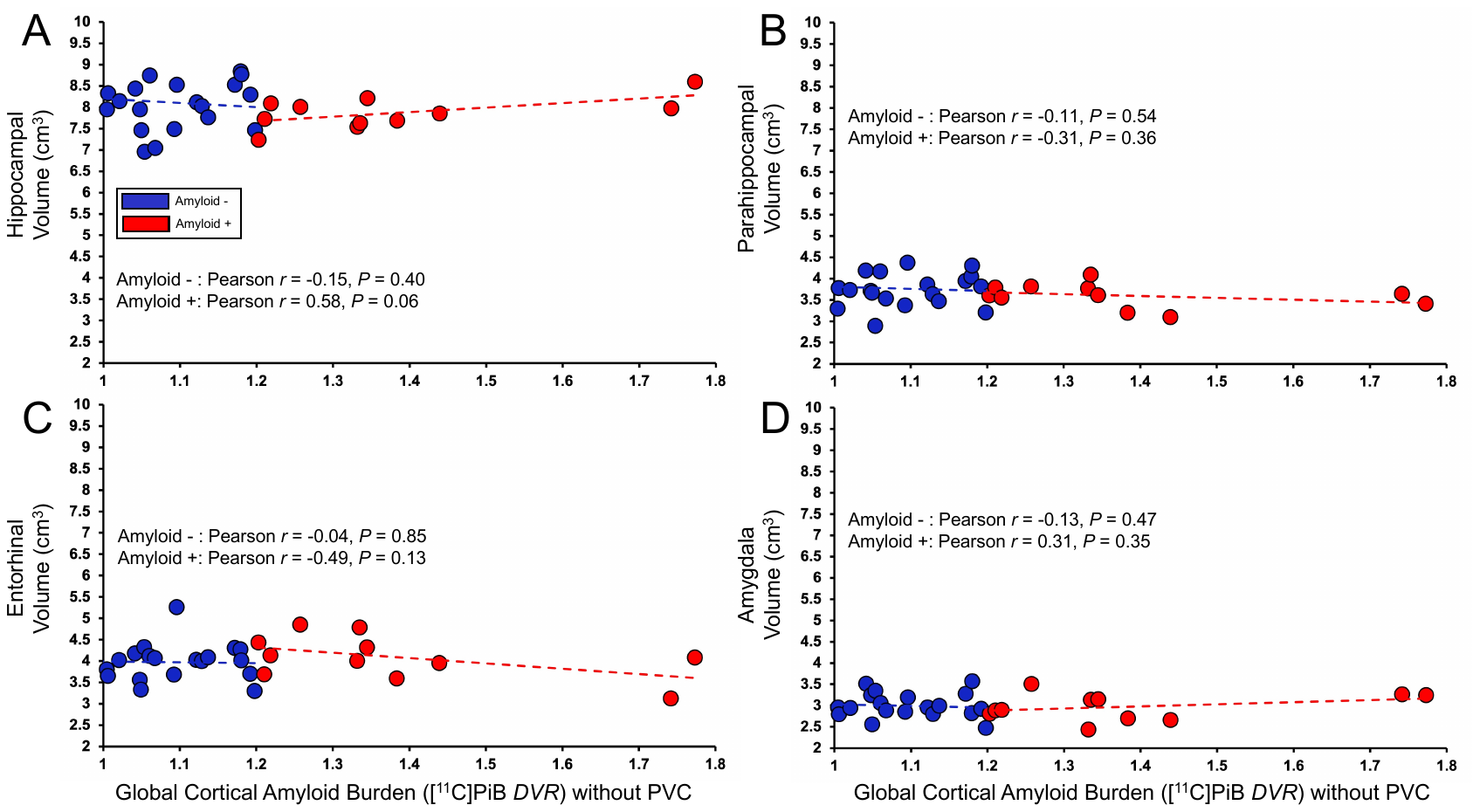

Association between mean global cortical amyloid burden and (A) hippocampal, (B) parahippocampal, (C) entorhinal, and (D) amygdalar volume in Aβ + and Aβ- participants using a [^11^C]PiB *DVR* threshold of 1.2. Data are Pearson *r* (*P*) for correlations between mean global cortical amyloid burden (*DVR*) and medial temporal lobe volumes, with separate models for Aβ + and Aβ- participants. A *DVR* threshold of 1.2 was used to determine amyloid status (positive vs. negative). Global cortical Aβ burden was calculated as a weighted average of composite regions commonly affected by Aβ burden in AD, which included prefrontal, lateral temporal, posterior cingulate/precuneus, and lateral parietal ROIs. Abbreviations: *Aβ*, amyloid beta; *AD*, Alzheimer’s disease; *PiB*; Pittsburg compound B; *DVR*, distribution volume ratio.

**Supplementary Figure 9.**

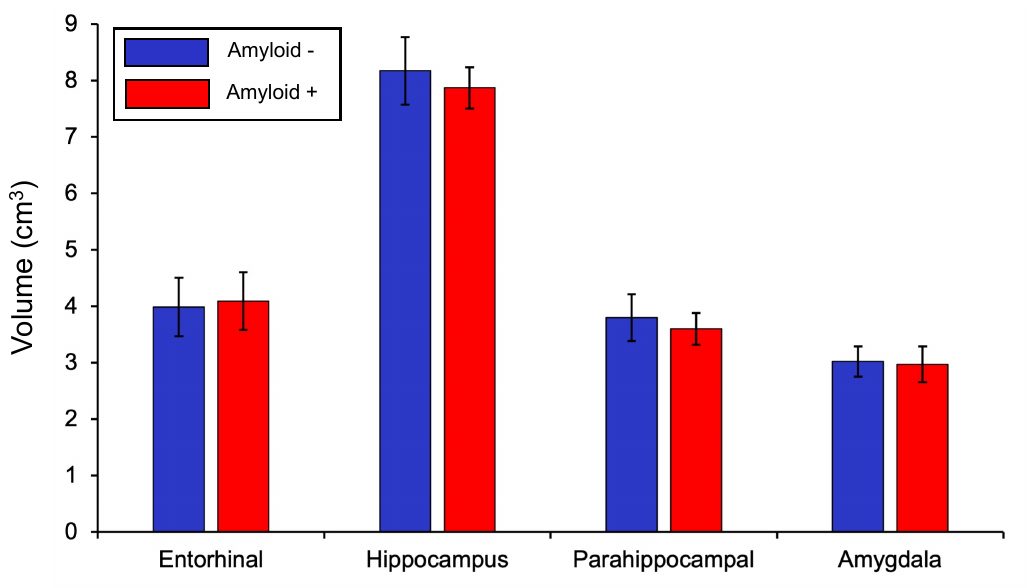

Group differences in medial temporal lobe volume by amyloid status using a mean global cortical amyloid *DVR* threshold of 1.2. Bar graphs represent mean (SD) for medial temporal lobe volumes (cm^3^) for amyloid positive and negative participants. Volumes were normalized using each participant’s eICV. Group differences were determined using unpaired t-tests and *P* values are depicted above the bar graphs to denote significant group differences. A mean global cortical [^11^C]PiB *DVR* threshold of 1.2 was used to determine amyloid status (positive vs. negative). Global cortical Aβ burden was calculated as a weighted average of composite regions commonly affected by Aβ burden in AD, which included prefrontal, lateral temporal, posterior cingulate/precuneus, and lateral parietal ROIs. Abbreviations: *Aβ*, amyloid beta; *AD*, Alzheimer’s disease; *PiB*; Pittsburg compound B; *DVR*, distribution volume ratio; *eICV*, estimated intracranial volume.

**Supplementary Figure 10.**

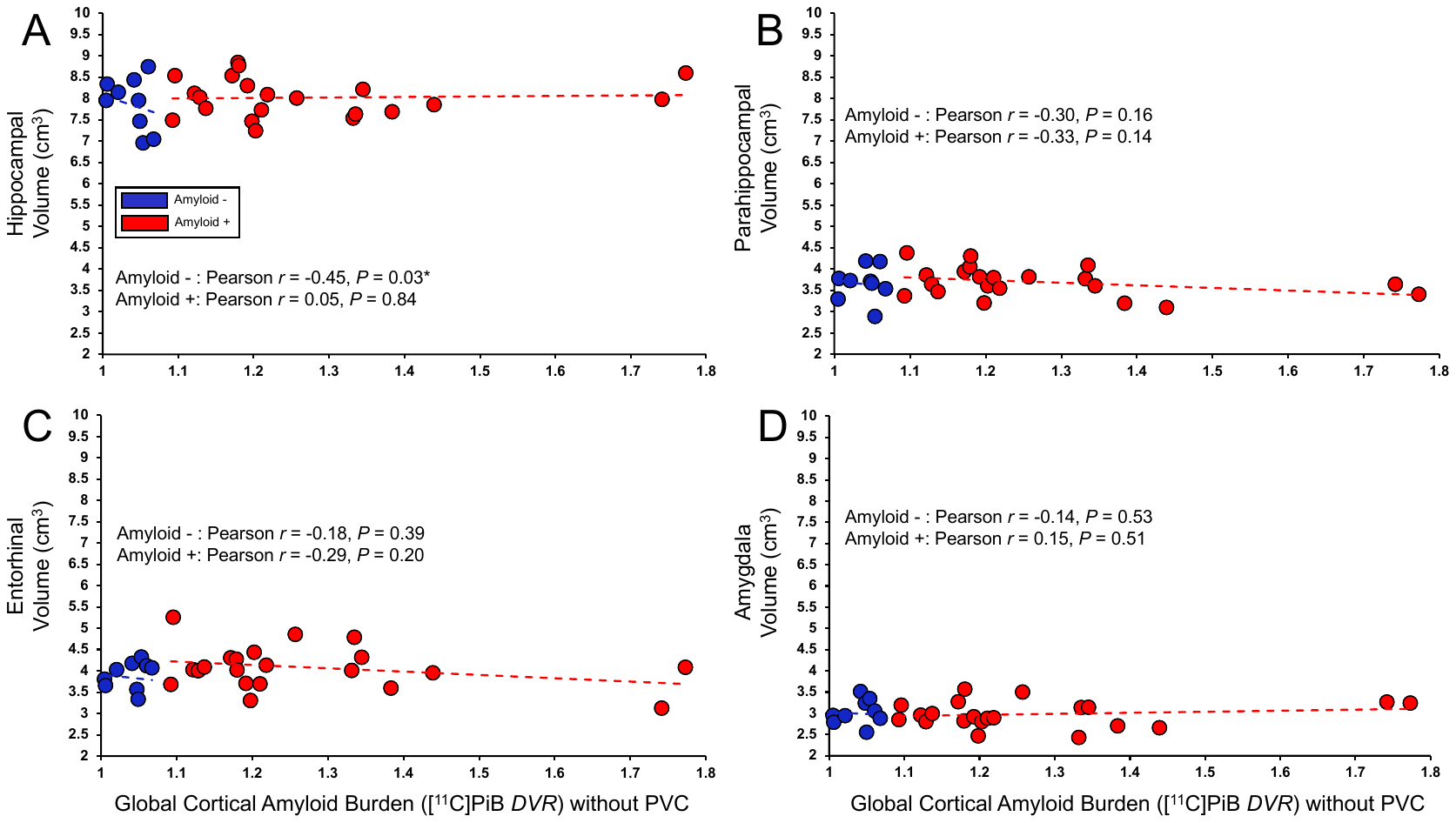

Association between mean global cortical amyloid burden and (A) hippocampal, (B) parahippocampal, (C) entorhinal, and (D) amygdalar volume in Aβ + and Aβ- participants using a [^11^C]PiB *DVR* threshold of 1.08. Data are Pearson *r* (*P*) for correlations between mean global cortical amyloid burden (*DVR*) and medial temporal lobe volumes, with separate models for Aβ + and Aβ- participants. A *DVR* threshold of 1.08 was used to determine amyloid status (positive vs. negative). Global cortical Aβ burden was calculated as a weighted average of composite regions commonly affected by Aβ burden in AD, which included prefrontal, lateral temporal, posterior cingulate/precuneus, and lateral parietal ROIs. Abbreviations: *Aβ*, amyloid beta; *AD*, Alzheimer’s disease; *PiB*; Pittsburg compound B; *DVR*, distribution volume ratio.
